# Transcriptomic analysis of pathways associated with αv integrin-related non-canonical autophagy in human B cells

**DOI:** 10.1101/2021.08.02.452710

**Authors:** Virginia Muir, Sara Sagadiev, Emmaline Suchland, Iana Meitlis, Natalia Giltiay, Jenny M. Tam, Ethan C. Garner, Carl N Wivagg, Donna Shows, Richard G. James, Adam Lacy-Hulbert, Mridu Acharya

## Abstract

Autophagy proteins have been linked with development of immune-mediated diseases including lupus, but the mechanisms for this are unclear. We have previously shown that non-canonical autophagy induced by αv-integrins regulates B cell activation by viral and self-antigens in mice. Here we investigated the involvement of this pathway in B cells from human tissue. Our data revealed that autophagy is specifically induced in germinal-center and memory B cell sub-populations from human tonsil and spleen. Transcriptomic analysis showed that induction of autophagy is related to unique aspects of activated B cells such as mitochondrial metabolism. To understand the function of non-canonical autophagy in B cells, we used CRISPR-mediated knockdown of autophagy genes. Integrating data from primary B cells and knockout cells we found that αv-integrin-related non-canonical autophagy limits activation of specific pathways while promoting others. These data provide new mechanistic links for autophagy and immune dysregulation in diseases such as lupus.

## Introduction

Conventionally, autophagy is an evolutionarily conserved intracellular process for recycling of organelles and degradation of unwanted cytoplasmic cargo. Canonical autophagy (known as macro-autophagy) is activated during cell stress or starvation and is involved in various fundamental cellular processes such as growth and differentiation (1, 2). During macro-autophagy interactions between a series of autophagy related proteins lead to lipidation of the ubiquitin-like protein LC3 and its recruitment to double membrane vesicles called autophagosomes, which deliver cytoplasmic contents for degradation or recycling to the lysosomes. Additionally, autophagy related proteins and LC3 have also been shown to be involved in other intracellular trafficking events related to signaling and processing of pathogen-derived materials that do not require the formation of the autophagosomes (3-5). These processes are referred to by different autophagy related terms for more non-canonical roles of autophagy proteins. Genome-wide association studies (GWAS) have implicated autophagy proteins in immune-mediated diseases such as inflammatory bowel diseases and Systemic lupus erythematosus (SLE), making them interesting targets for modulation of immune response (6-8).

In B cells of the immune system, autophagy has been shown to be involved in differentiation of the B1-B cell subset, antibody production by plasma cells and for maintenance of memory B cells (9-12). More recently non-canonical autophagy has also been shown to be activated in B cells (13). However, the exact functions of non-canonical autophagy in B cells and its mechanism of activation have remained unclear. We have previously shown that signaling through the αv family of integrins can activate autophagy proteins which then regulate the rate and duration of toll-like receptor (TLR) signaling in B cells (14). Specifically, we have found that upon B cell stimulation by TLR ligands, αv integrins are activated, leading to activation of Src and Syk kinases. Syk activation then leads to activation of the autophagy proteins and LC3 lipidation, which functions as a key regulator of outcome of signaling. The central feature of this regulation is that LC3 lipidation is required for the TLRs to transition from NF-κB signaling endosomes to IRF7 signaling late endosomes. LC3 activation also eventually leads to termination of TLR signaling in the late endosomal compartments. Loss of αv integrins or autophagy proteins leads to altered TLR trafficking and signaling and particularly the loss of autophagy proteins leads to absence of IRF7 signaling; Overall these results indicated that autophagy proteins while curtailing NF-κB signaling promote IRF7 signaling. We further showed that this type of fine-tuning of TLR signaling is important in regulating B cell response to viral antigens as well as in regulating self-reactive B cell responses during development of lupus-like disease in mice (15, 16). This role of autophagy proteins in B cells does not require formation of the double membraned autophagosomes hence it is a form of non-canonical autophagy. We have termed it as TLR induced autophagy (TIA) or activation induced autophagy related to B cell activation by TLR or BCR ligands. Furthermore, other groups have also shown the importance of autophagy proteins in regulating TLR signaling in macrophages and plasmacytoid dendritic cells (17, 18).

This function of autophagy proteins in fine-tuning the rate and duration of TLR signaling and coordinating signaling through other cell surface receptors such as the integrins begins to shed light on how these proteins might be involved in regulating immune activation in diseases such as SLE. Based on our mouse studies, we were keen to assess the role of these proteins in B cell activation during lupus. But before assessing the status of this pathway in B cells from a disease micro-environment, it was essential to establish how these proteins are functioning in human B cell populations. B cells from the bone marrow enter circulation as transitional B cells which differentiate into mature naïve B cells. Naïve B cells encounter antigens through their B-cell receptor (BCR) while other receptors such as TLRs also get activated during antigenic stimulation. This leads to generation of more activated mature B cell subsets such as germinal center (GC) B cells, memory B cells, and plasma cells. These activated mature B cell subsets each have varying capacity to respond to antigenic stimulation and inherent differences in their signaling capacities ensure the most appropriate B cell response to antigens. In order to get an overall picture of the role of autophagy proteins in B cell subsets, we examined B cells from human tonsil tissue, which allows us to isolate a number of different mature B cell populations (transitional, naïve, GC and memory B cells) (19). Analysis of activation of autophagy pathway in these subsets showed striking differences in induction of this pathway in each B cell subsets leading to the questions why this was the case and how this relates to the function of these proteins in B cells.

We hypothesized that the autophagy pathway is upregulated in certain B cell populations because autophagy proteins have important roles in activated B cells both through macro-autophagy and non-canonical autophagy like TIA. While macro-autophagy could play important role in regulating functions such as protein secretion by plasma cells, we predict non-canonical autophagy plays a role in adjusting responses of activated B cells. Alterations in these types of adjustment mechanisms could lead to the aberrant activation of B cells seen in diseases such as lupus. To understand these functions of autophagy proteins in more detail, we leveraged the differences in the induction of autophagy pathway in B cell subsets to identify genes and pathways related to autophagy proteins. We then combined this with the analysis of genes and pathways specifically regulated by non-canonical autophagy in an activated B cell line to define the consequences of activation of this pathway in human B cells.

## Results

### B cell subsets from human tissues show distinct differences in activation of autophagy pathway

We have previously shown that TLR-Induced autophagy (TIA) preferentially occurs in activated B cell subsets in mice, including B-1, MZ and GC B cells. To determine whether human B cells show similar differences, we isolated transitional, naïve, GC and memory B cells from tonsils (**Fig 1A**) and measured activation of autophagy following TLR stimulation. We assessed autophagy by measuring the appearance of lipidated LC3 (LC3-II) by immunoblot (14, 16, 20), in response to treatment with the ligands for TLR7 and TLR9. Cells were also treated with Rapamycin, which induces macro-autophagy, to determine the cells capacity for LC3-lipidation, and with Chloroquine, which blocks endo-lysosomal acidification and degradation of LC3, allowing measurement of basal autophagy activity in cells.

**Fig 1:**
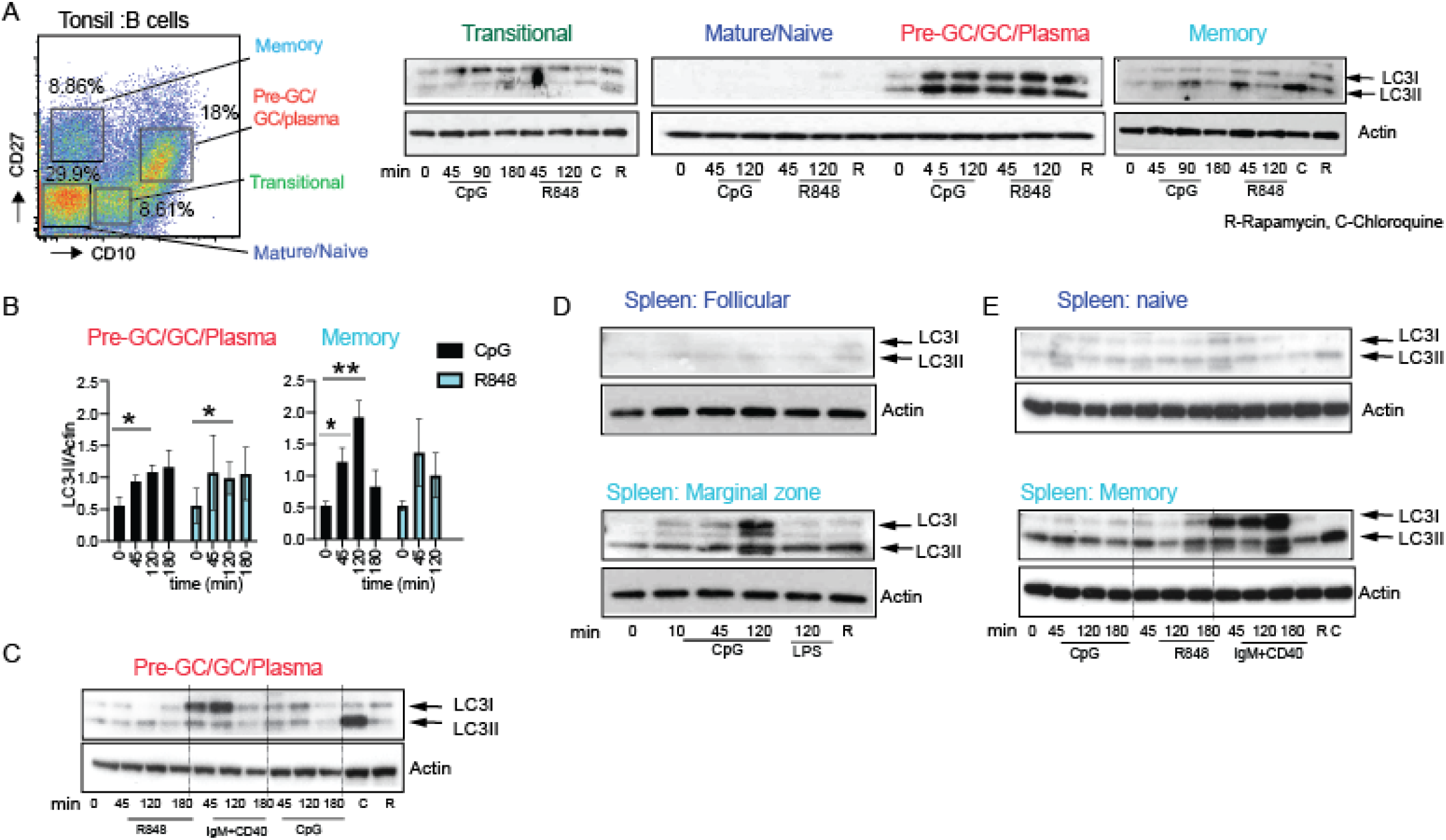
Autophagy proteins are specifically induced in activated B cell subsets from human tissue: **(A)** Flow cytometry gating for sorting the different B cell subsets (CD19+) and western blot analysis for LC3 expression and processing in sorted B cell subsets (200,000-250,000) from human tonsils is shown. B cell subsets were sorted based on CD19, CD10 and CD27 expression and additional analysis of these subsets was used to delineate them as: naïve B cells (CD19+ CD10 low CD27 low, IgM ++, IgD+), transitional B cells (CD19+, CD10++, CD27 low, IgM++, IgD+), memory B cells (CD19+, CD10low CD27++, IgM ++, IgD-) and preGC/GC, plasma cells(CD19+, CD10++++, CD27++, IgM+/-, IgD-). Sorted B cells were stimulated for the indicated times with CpG DNA (3mM) or R848 (5mg/ml) or Rapamycin(R) (0.6mM) or chloroquine (0.1mM) (C) for indicated time (mins) and cytoplasmic extracts from cell lysates were used to stain for LC3. Actin is shown as the loading control. **(B)** Quantification of LC3 lipidation as measured by changes in the LC3-II band and Actin by densitometry. **(C)** LC3 processing upon stimulation with anti-IgM (10μg/mL) and CD40 ligand (CD40L) (5μg/mL). **(D-E)** Analysis of LC3 processing in B cell subsets Marginal zone (CD19+ CD27+ CD23neg), follicular (CD19+, CD27neg CD23+) naïve (CD19+ CD10neg CD27neg) and memory (CD19+ CD10neg CD27+), sorted from human spleens. Data are representative blots based on blots from 8 different tonsil donor samples and 5 different spleen donor samples. Quantification of data from 3-5 individual donors are shown, and significance indicated as by * (*p*<0.05) or ** (*p*<0.01). *p*-values calculated using Mann Whitney test.

Immunoblots revealed distinct patterns of LC3 expression and lipidation in each subset. Naïve B cells expressed lowest amount of either LC3-I or LC3-II compared to other subsets (**Fig 1A)**, and in all the donors tested we did not observe a strong induction of LC3-II after treatment with TLR ligands or Rapamycin. Transitional B cells expressed primarily LC3-I. Treatment with Rapamycin and Chloroquine caused accumulation of low levels of LC3-II, indicating these cells undergo autophagy, but LC3-II was not generated following TLR stimulation (**Fig 1A**). Memory B cells had low levels of LC3-I at baseline and showed a robust induction of LC3-II after TLR stimulation and treatment with Rapamycin or Chloroquine (**Fig 1A-B**). GC B cells had the highest expression of LC3-I, as well as high basal levels of LC3-II in many donors. The intra-donor variation probably reflects the heterogeneity of this population, which includes pre-GC cells and some plasma cells. LC3-II levels increased after TLR or Rapamycin stimulation in the GC cells **(Fig 1A-B)**, but LC3-II induction upon stimulation was less evident in the donors with high basal autophagy. BCR/CD40 stimulation also led to LC3-II induction and the major effect of this stimulation was on increasing LC3-I (**Fig 1C**). While TLR stimulation also led to some increase in LC3-I expression, overall our data indicate that TLR stimulation promotes lipidation of LC3-I to LC3-II but BCR/CD40 might have a major effect on LC3-I.

To confirm and extend these findings, we analyzed LC3 expression in human splenic B cells. Naïve (CD27neg CD10 neg) and Follicular (CD27neg CD23+) spleen B cells showed very little expression of LC3-I or induction of autophagy. In contrast, memory B cells and marginal zone-like B cells had higher levels of LC3-I, and generated LC3-II after TLR stimulation (**Fig 1D,E**). As for tonsil GC B cells, spleen memory B cells also generated LC3-II after BCR/ CD40 stimulation with major effect on LC3-I (**Fig 1E**). Thus, based on these data, we concluded that ‘activated’ B cells, including memory, GC and marginal zone-like B cells are the principal B cells that can undergo autophagy, and do so in response to both TLR stimulation and activation through the BCR/CD40.

TIA is required for activation of the transcription factor IRF7 downstream of TLR7 and TLR9 (14, 16, 18). To determine whether TIA corresponded with IRF7 activation in human B cells, we measured nuclear NF-

κB and IRF7 levels in tonsil and spleen B cell subsets after stimulation. We quantified changes in the kinetics of activation of these transcription factors in subsets where we had cells for multiple donors (**Sup Fig 1C**). NF-κB was activated by TLR and BCR/CD40 stimulation in all cells tested. IRF7 activation, however, was only seen at high levels in tonsil GC and memory cells, and in spleen marginal zone-like cells (**Sup Fig 1**). Although low levels of activation were seen in mature/ naïve B cells these were not consistent among donors. Hence, as we have seen in mouse B cells, activation of autophagy corresponds with a switch from NF-κB to IRF7 signaling in activated B cell subsets.

### av-signaling is activated in GC and memory B cells subsets

We have previously shown that TIA in mouse B cells requires αv integrins and involves αvβ3 integrin activation and internalization. We therefore next determined whether changes in αv integrin expression or activation were associated with TIA in human B cell subsets. By flow cytometry, we observed that αvβ3 and the related heterodimer αvβ5 were uniformly expressed by all tonsil B cells with no major differences in surface expression between cell subsets (**Fig 2A**). Integrin activity is often regulated by associated proteins, including members of the tetraspanin family, which promote activation and internalization of integrins, and have been associated with non-canonical autophagy (21). Notably, the tetraspanin CD9, which forms a complex with αv integrins (22), was upregulated in tonsil GC B cells, suggesting that GC B cells may be primed for activation of αvβ3. Likewise, the tetraspanin CD82, which also binds integrins, was upregulated on tonsil memory B cells (**Fig 2B**).

**Fig 2:**
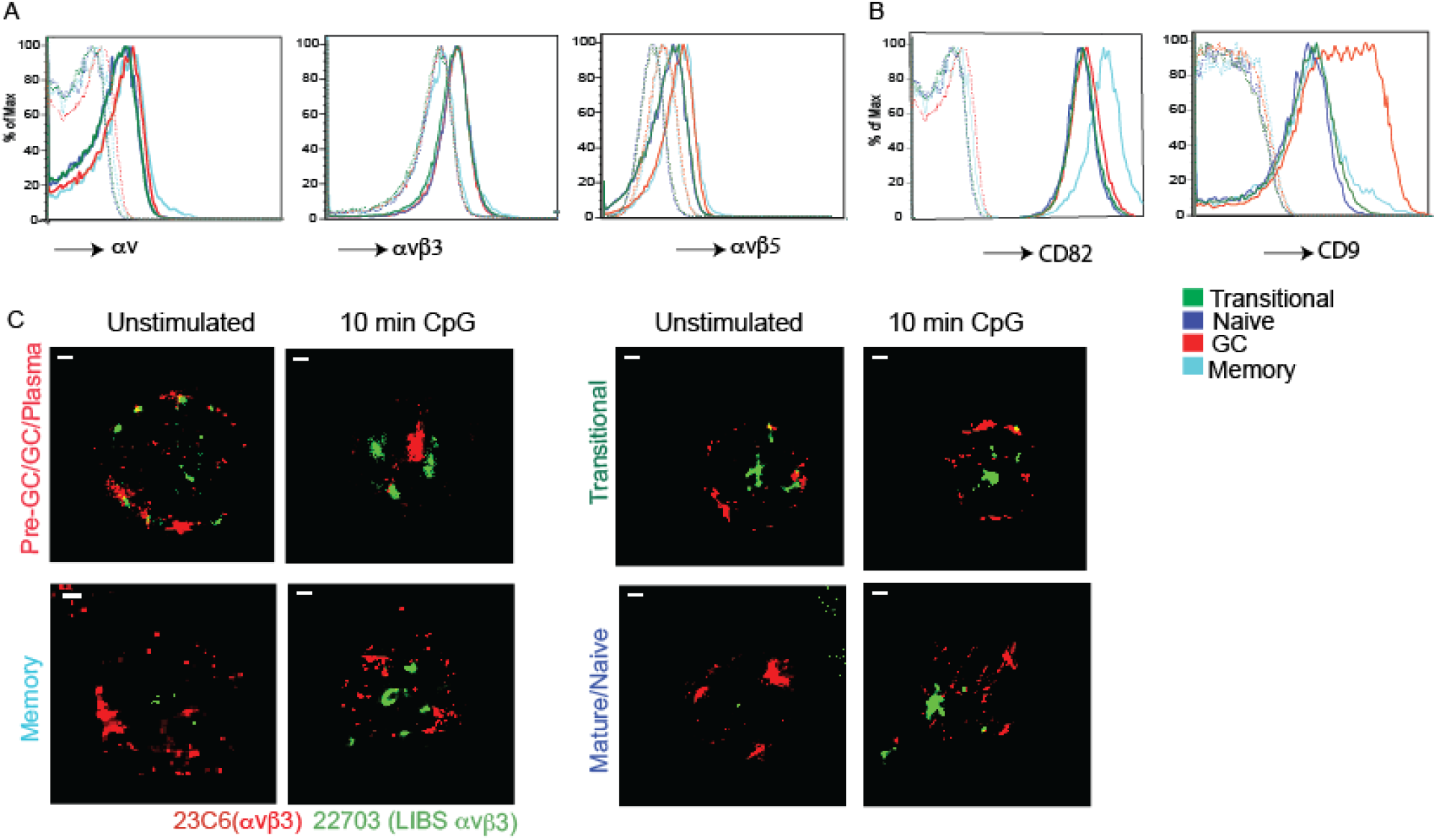
αv integrin expression and activation in tonsil B cell subsets: **(A-B)** Flow cytometry for staining of integrin heterodimers **(A)** or tetraspanins CD9 and CD82 **(B)** on the tonsil B cell subsets. Tonsil B cell subsets were distinguished as in **Fig 1A**. Dotted lines indicate cells without the specific integrin heterodimer or tetraspanin stain or cells stained with streptavidin APC alone. **(C)** STORM images for staining of sorted tonsil B cell subsets with anti-αvβ3 antibody and anti-αvβ3 LIBS antibody with or without CPG stimulation.

To test whether TIA in GC B cells was associated with integrin activation, we used microscopy to analyze integrin localization and activation in sorted tonsil B cell subsets. We turned to super-resolution microscopy for these studies which provides better resolution of integrin localization compared to conventional confocal microscopy. Using 23C6 antibody, which recognizes the αvβ3 heterodimer, we observed that this integrin was localized at the outer membrane of all cell subsets with little difference in the overall level or pattern of staining between subsets, consistent with FACS analysis. Following stimulation of GC cells with CpG DNA, αvβ3 re-organized to large intracellular compartments, as seen in our previous report of endo-lysosomal αvβ3 localization in TLR-stimulated MZ and B-1 B cells (14). Moreover, stimulation led to aggregates of αvβ3 that stained strongly with another αvβ3 antibody, 22703, which specifically recognizes the ligand induced binding site (LIBS) on activated αvβ3 (23). CpG treatment of both memory and naïve B cells led to αvβ3-LIBS staining, indicating that TLR signaling activated αvβ3 in these cells. But αvβ3 aggregates staining with LIBS antibody were most prominent and similar in GC and memory B cells compared to naïve cells. In contrast, CpG treatment had little effect on αvβ3 activation or localization in transitional B cells. We concluded that αvβ3 activation and internalization to large aggregates correlates with induction of autophagy in human GC and memory B cells, consistent with our previous observations in mouse MZ and GC B cells.

### Transcriptional analysis identifies a role for mitochondria-related genes in activation of autophagy proteins in GC B cells

To investigate genes and pathways related to activation of autophagy in the B cell subsets, we identified genes differentially expressed between sorted tonsil B cell subsets using RNA-Sequencing (RNA-Seq) (**Fig 3A**). Naïve and transitional B cells showed very similar patterns of gene expression, with few differences between these subsets. Memory B cells had similar expression patterns to naïve cells, with some distinct sets of differentially expressed genes. GC B cells showed the most distinct differences in gene expression from other subsets. We therefore focused on comparisons of gene sets and functional modules (24-26) between GC B cells, which undergo autophagy and naïve B cells, which do not show high levels of basal or TLR-induced autophagy.

**Fig 3:**
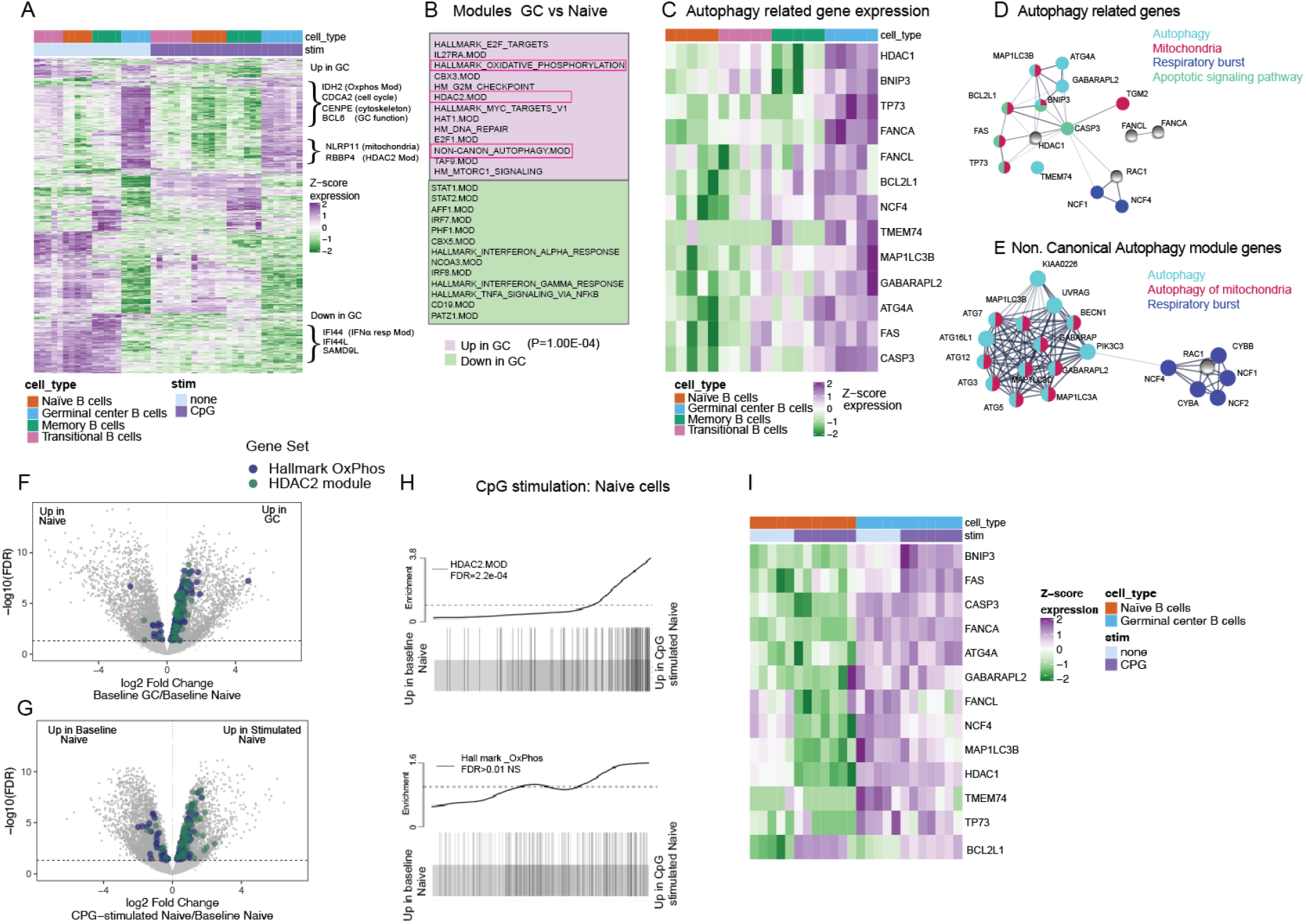
Expression and activation of autophagy pathway in activated B cells is related to upregulation of mitochondrial proteins: Sorted B cell subsets with or without stimulation (3h) with CpG DNA were used for RNA-sequencing. Cells were stimulated with either CpG 2395 or CpG 2006 and data includes results from both stimulations. **(A)** Gene expression heat map generated using the 350 most significantly DE genes from each subset compared to other subsets. Genes were hierarchically clustered using Euclidean distance & complete linkage; colors represent scaled gene expression values. Genes upregulated or downregulated in GC cells are highlighted. **(B)** Modules up-regulated or down-regulated in GC cells compared to Naïve cells, based on Roast analysis. 13 modules from each subset are shown and P-value from mroast analysis is indicated. (**C)** Heat map of selected autophagy genes’ scaled expression in the different B cell subsets. **(D)** String network analysis for gene-gene connection among autophagy related genes’ that are enriched in GC cells. Genes are presented as circular nodes and known gene to gene interaction from STRING are shown as gray lines. Thickness of the line indicates strength of data support. **(E)** String network analysis of the genes from the Non.Canonical Autophagy module. **(F-G)** Volcano plots showing differentially expressed genes in Naive cells compared to GC cells or Naïve cells with and without CpG stimulation. Highlighted are the genes belonging to Oxidative phosphorylation and HDAC2 modules from gene set enrichment analysis. **(H)** Barcode plots showing enrichment of HDAC2 or Hallmark Oxphos module genes in Naïve cells upon CpG stimulation. FDR (false discovery rate) is indicated in the plot. **(I)** Heat map of selected autophagy genes’ scaled expression in GC and Naïve B cell subsets with or without CpG stimulation. Data are based on RNA-sequencing from 5 different donor samples.

Many of the genes expressed at higher levels in GC cells reflect known differences in their functional properties, including genes involved in cell proliferation, metabolism, cytoskeleton and DNA repair (**Fig 3A**). Analysis of previously described gene modules involved in autophagy identified only one module, associated with Non.canonical autophagy (4) as significantly upregulated in GC B cells compared with naïve cells (**Fig 3B**). A deeper analysis of autophagy gene modules however, identified several autophagy-related genes whose expression correlated with autophagy-activity across B cell subsets, being low in naïve B cells, higher in transitional and memory B cells, and highest in GC B cells (**Fig 3C**). Most of the autophagy-related genes showing this pattern have previously been implicated in a specific form of autophagy, termed mitophagy, mitochondrial function or in ROS (Reactive oxygen species) production (**Fig 3D**) (27-30). Similarly, genes in the Non.canonical autophagy module up regulated in GC cells were also related to mitochondrial function or respiratory burst (**Fig 3E**). Furthermore, gene modules related to mitochondrial metabolism, including oxidative phosphorylation (Oxphos) and HDAC signaling, were also significantly elevated in GC B cells and in memory B cells (data not shown) compared to naïve cells (**Fig 3B and F**). These pathways were of particular interest as they have been linked to induction of autophagy and to generation of ROS which we have previously shown are required for TIA in mouse B cells (27, 28). While both these modules showed a trend towards up-regulation upon stimulation in naïve B cells, (**Fig 3G, H**) the autophagy related genes themselves did not show up-regulation upon stimulation in the naïve cells (**Fig 3I**). Hence, these data suggest that the activity of the autophagy pathway in B cell subsets is associated with expression of genes involved in mitophagy and mitochondrial metabolism and the activation of the autophagy pathway appears to be a feature acquired by GC B cells over time and not induced by short-term activation.

### Mitochondria-related genes are involved in TLR-induced autophagy in B cells

To confirm the role of mitophagy related genes in activation-induced autophagy in human B cells, we turned to a genetically tractable system, the B cell line, HBL1 (31). These cells resemble activated B cell stage and respond strongly to TLR9 stimulation. HBL1 cells expressed high levels of both LC3-I and LC3-II at baseline and showed robust accumulation of LC3-II after treatment with Chloroquine, indicating ongoing autophagy in these cells, probably due to their continual growth and proliferation in culture (**Fig 4A**). Despite this basal autophagy activity, stimulation with the TLR9 ligand CpG DNA (either CpG 2006 or CpG 2395) promoted further accumulation of LC3-II (**Fig 4A**). LC3-II accumulation was completely blocked by CRISPR deletion of the core LC3 conjugation components Atg5 and Atg7, which are essential for all forms of autophagy, confirming that our assay was measuring LC3-lipidation and that HBL-1 cells were undergoing autophagy (**Fig 4A**). We next tested whether activation-induced autophagy in HBL-1 cells required αvβ3 integrin. CRISPR-deletion of all αv integrins blocked LC3 lipidation induced by CpG DNA treatment but had no effect LC3-II accumulation after Chloroquine treatment. (**Fig 4B and C**) These data are consistent with our hypothesis that TLR stimulation promotes LC3 lipidation through an αv-dependent non-canonical autophagy pathway, distinct from the basal autophagy mechanisms.

**Fig 4:**
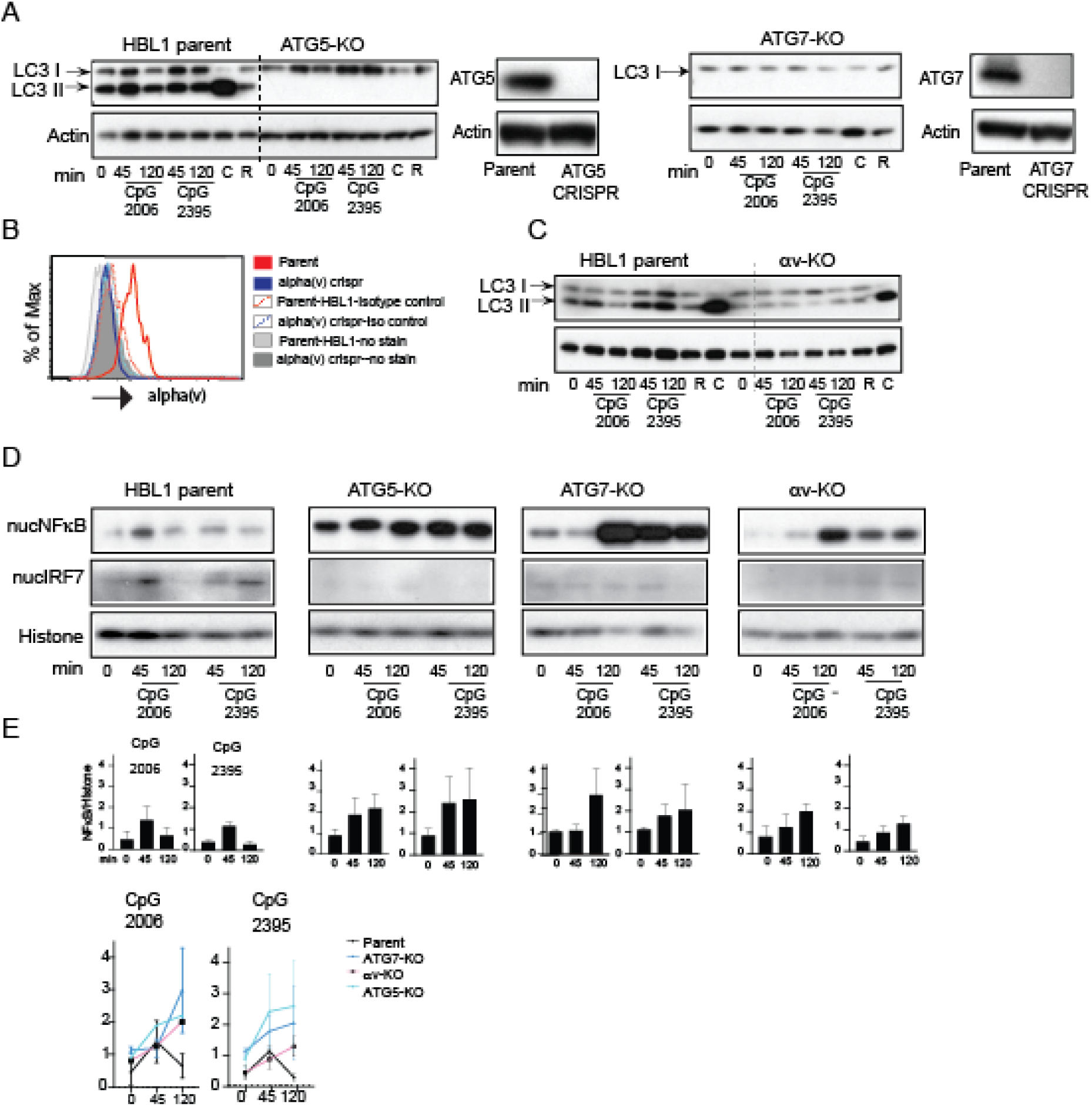
Non-canonical autophagy regulates TLR signaling in activated human B cell line: **(A)** Western blot analysis of LC3 processing or LC3 lipidation in parental HBL1 cells or cells with CRISPR of autophagy proteins (ATG5 or ATG7). Parental cells or CRISPR cells were stimulated with either CpG 2006, CpG 2395 or with autophagy modulators Rapamycin (R) or Chloroquine (C) for indicated times and cytoplasmic extracts were used to assess LC3. Actin is shown as loading control. Also shown are western blot analysis of changes in the protein levels of ATG5 or ATG7 in the CRISPR cells compared to parental cells. **(B)** Flow cytometry analysis for changes in protein levels of av integrin after CRISPR of the av gene. **(C)** LC3 lipidation in parental HBL1 cells or HBL1 cells CRISPR of av integrin **(D)** Nuclear fraction from the stimulated parental cells or cells with CRISPR of ATG5 or ATG7 or av integrins were analyzed for NF-kB and IRF7 activation by western blot. Histone is shown as nuclear loading control **(E)** Graphs showing changes in NF-kB upon either CpG2006 or CpG 2395 stimulation are quantified based on densitometry from at least 3 independent experiments. In all cases, representative blots from at least 3 independent experiments are shown.

Disrupting autophagy also affected TLR signaling in HBL-1 B cells (**Fig 4D and E**). In parental HBL1 cells, treatment with CpG caused rapid activation of NF-κB, which returned to basal levels by 120 minutes, followed by activation of IRF7, similar to our observations in tonsil GC B cells. Both *ATG5*-knockout and *ATG7*-knockout HBL-1 cells had increased basal levels of NF-κB and increased and sustained NF-κB activation after CpG stimulation. In contrast, IRF7 was not activated after CpG treatment in autophagy-knockout cells. Knockout of *ITGAV* had similar effects to disruption of all autophagy, causing increased and sustained activation of NF-κB, and loss of IRF7 activation **(Fig 4D and E**).

To ask whether the increase in mitophagy genes in the GC B cells was related to activation-induced autophagy, we disrupted the Bcl-family member BNIP3. BNIP3 is critical for initiation of mitophagy in several cell types (30, 32) and we found it to be expressed at higher levels in GC B cells compared to naïve cells (**Fig 3C**). *BNIP3*-knockout HBL-1 cells decreased LC3 lipidation after stimulation with CpG DNA (**Fig 5A and B)**. As we had observed for *ITGAV*-knockout cells, disruption of BNIP3 did not affect LC3-II levels in unstimulated cells or following Chloroquine treatment. Similar to *ITGAV*-knockout, *BNIP3* knockout also resulted in sustained NF-κB activation and loss of IRF7 activation. Hence, both αv integrins and BNIP3 are required for activation-induced autophagy in human B cells, indicating that this occurs through a mitophagy-like process, but are not required for ‘basal’ autophagy, which likely therefore occurs through a separate pathway, such as macro-autophagy. Thus, these data demonstrate a role for mitophagy genes in activation-induced autophagy in B cells and confirm that this process curtails NF-κB signaling downstream of TLR activation but is required for activation of IRF7.

**Fig 5:**
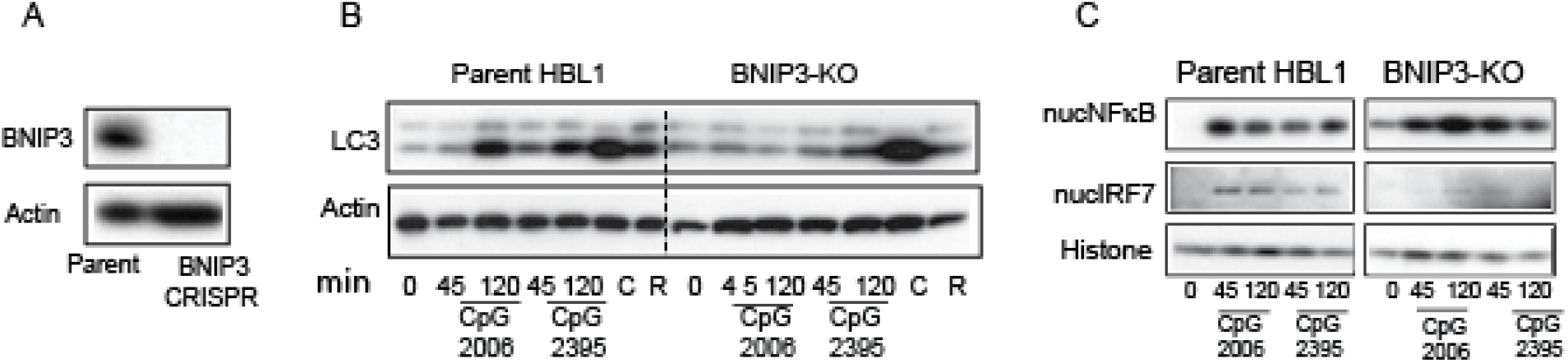
BNIP3 is involved in regulating TLR induced autophagy in human B cells: **(A)** Western blot analysis shows changes in the protein levels for BNIP3 in the CRISPR cells compared to parental cells. **(B)** Western blot analysis of LC3 processing in parental HBL1 cells or cells with CRISPR of the mitochondrial protein BNIP3 Parental cells or CRISPR cells were stimulated with CpG or with autophagy modulators Rapamycin (R) or Chloroquine (C) for indicated times and cytoplasmic extracts were used to assess LC3. Actin is shown as loading control. **(C)** Nuclear fraction of the stimulated cells was analyzed for NF-κB and IRF7 activation by western blot. Histone is shown as nuclear loading control. Representative blots from at least 3 independent experiments are shown.

### Autophagy regulates transcriptional response to stimulation in GC B cells

We reasoned that the effects of activation-induced autophagy on signaling would impact transcriptional responses in B cells. To investigate this, we analyzed gene expression in tonsil B cell subsets after stimulation through TLRs or BCR, focusing on naïve and GC B cells, as these subsets had the clearest differences in induction of autophagy. Naïve B cells showed extensive gene expression changes after stimulation, with over 6500 differentially expressed genes across all three conditions (CpG DNA, R848, or combined anti-IgM and CD40-ligand). The majority of these genes were similarly altered in response to all stimuli, although we also identified genes that were specific to each stimulation. Fewer genes were affected by stimulation in GC B cells (4423 genes differentially expressed across all conditions compared with 6717 genes in naïve cells) and responses to stimulation were similar for TLRs and BCR/CD40 in GC cells, unlike what we observed in naïve B cells **(Sup Fig2A**).

To compare responses between GC and naïve B cells, we first focused on changes in pre-defined and curated gene sets or modules, based on common function or co-expression patterns. Majority of the modules differentially expressed after stimulation in GC cells were also altered in stimulated naïve cells. Additional gene modules were differentially expressed after stimulation in naïve B cells but not in GC cells, reflecting the higher numbers of differentially expressed genes in naïve cells, (**Sup Table 1)**. In many cases, GC B cells showed smaller changes in expression of modules after stimulation compared to naïve cells **(Fig 6A and Supp Fig 2**), and this was often associated with changes in basal expression between the two cell subsets. As examples, three modules upregulated by stimulation in both subsets (Myc target version 2, HDAC2 and Unfolded Protein Response) had higher basal expression in GC cells but were less strongly upregulation after stimulation (**Fig 6A**). Similar patterns were seen for gene modules downregulated after stimulation in GC and naïve cells. We did not identify any modules that were significantly upregulated by stimulation only in GC cells, but two modules related to cell cycle and proliferation (Hallmark G2M checkpoint and E2F) were significantly downregulated by all stimuli in GC cells only. As autophagy is associated with changes in NF-κB and IRF7 activation, we analyzed expression of gene modules connected with these transcription factor pathways. NF-κB-responsive genes showed reduced response in stimulated GC B cells compared with naïve B cells (**Fig 6B**). For IRF7, one of the modules tested which is based on genes identified from a B cell line (33) showed a trend towards upregulation upon stimulation in the GC cells **(Fig 6B)**. This is in agreement with our hypothesis that autophagy limits induction of NF-κB related genes but promotes expression of IRF7 related genes.

**Fig 6:**
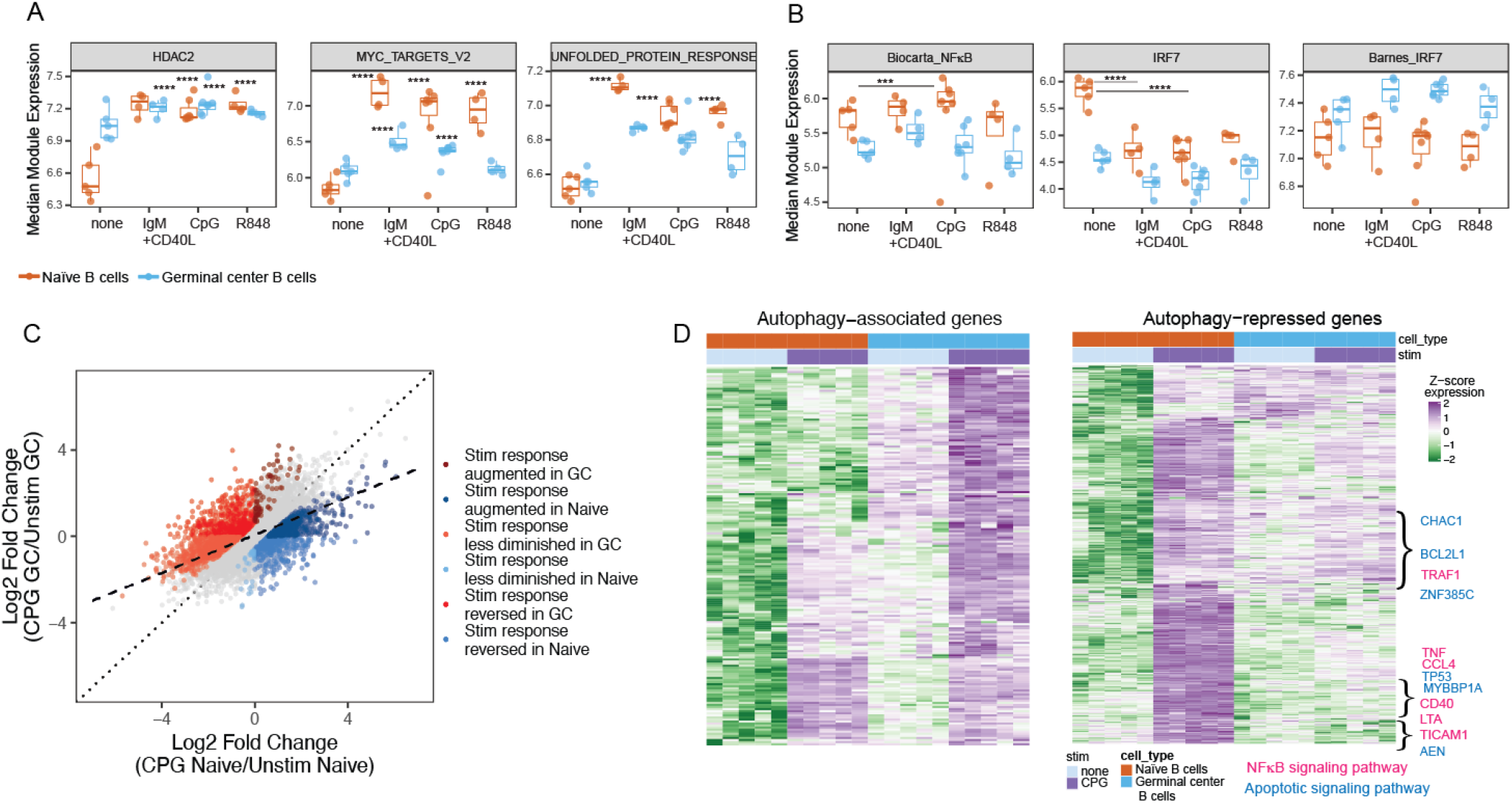
Autophagy proteins limit TLR responses in activated human B cells: Sorted B cells with or without stimulation (3h) with either CpG or R848 or anti-IgM +CD40L were used for RNA-sequencing. Data for CpG includes both cells stimulated with CpG 2395 or with CpG2006. **(A)** Graphs show as log2 TMM-normalized median module expression of gene sets in Naïve and GC cells using ROAST analysis. Box indicates interquartile range. **(B)** Median module expression for NF-κB and IRF7 in GC vs naïve cells with or without stimulation. p-values from mroast gene set tests are indicated as ***(*p*<0.001) or ****(*p*=1×10^−4^). **(C)** Graph showing gene expression changes upon CpG stimulation in GC cells compared to naïve cells. Dotted line represents perfect correspondence between CpG response in Naïve and GC cells. Dashed line is fit to observed data. Non-grey dots highlight genes exhibiting significantly different stimulation responses between Naïve and GC cells: In dark red are genes which increase to a greater degree in GC cells upon stimulation than in Naïve cells and in dark blue are genes which increase to a greater degree in Naïve cells than in GC cells. In light red are genes that decrease to a greater degree in Naïve cells upon stimulation than in GC cells and in light blue are genes that decrease to a greater degree in GC cells than Naïve cells. In medium red are genes that show increased expression in GC cells upon stimulation but show decrease in expression in Naïve cells and in medium blue are genes that show increase in expression in Naïve cells upon stimulation but show the opposite response in GC cells. **(D)** Heat map of genes showing increased basal increase in GC cells and induced upon CpG stimulation (GC_0 > N_0 & GC_CPG > GC_0) is shown as the autophagy associated set and heat map of genes induced upon CpG stimulation in the Naïve cells that show increased stimulation response in the naïve cells compared to GC cells (N_CPG > N_0 & CPG_resp_N > CPG_resp_GC) is shown as the autophagy repressed set. Genes were hierarchically clustered using Euclidean distance & complete linkage; colors represent scaled gene expression values. Data are derived from RNA-sequencing on samples from 5 different donors.

To determine whether the changes observed in gene module analysis were also seen at the level of individual genes, we next focused on transcriptional responses to CpG stimulation by GC and naïve B cells. We identified 2475 genes with differential responses to stimulation in the two cell subsets, (represented by all of the colored dots) (**Fig 6C**). Of the genes that were upregulated in response to stimulation in both cell subsets, the majority (586/ 664) showed a stronger response in naïve cells (dark blue dots), whereas only 78 showed stronger induction in GC cells (dark red dots). A similar pattern was seen for downregulated genes, with the majority showing stronger responses in naïve (coral dots) than GC cells (light blue dots) (**Fig 6C**).

Taken together, these analyses of genes and modules indicated that GC B cells generally showed smaller changes in gene expression after stimulation than naïve cells. For a subset of genes, this was associated with higher levels of basal gene expression in GC cells, consistent with their activated phenotype. We predict that non-canonical autophagy plays a role in this moderate response of GC cells and we used these data to generate a list of genes that showed moderate response in GC cells compared to naïve cells. We hypothesized that this list would contain potential *autophagy repressed* genes (**Fig 6D**). We further identified a smaller set of genes that were preferentially upregulated after stimulation in GC cells, which may represent genes whose expression is promoted by autophagy (*autophagy associated* genes) (**Fig 6D)**. String pathway analysis showed that while both these gene sets included genes associated with mitochondrion, metabolism and exosomes the *autophagy repressed* set included additional genes associated with the NF-κB signaling and apoptotic signaling pathway (**Fig 6D**).

### Non-canonical autophagy limits specific pathway while promoting others

To confirm the role of non-canonical autophagy in regulating genes or pathways identified from primary B cells, we made use of the HBL-1 cells in which we could disrupt autophagy by gene deletion. HBL1 cells with deletions in the core autophagy components *ATG5* and *ATG7*, or in *ITGA*V, were left unstimulated or stimulated with CpG DNA and gene expression analyzed by RNA-Seq (**Fig 4**). Knockout cells showed differential gene expression patterns from unmanipulated HBL1 cells (**Sup Fig 3A**). Furthermore, *ATG7* and *ITGAV*-knockout cells showed similar patterns of gene expression while the *ATG5-*knockout appeared to be more distinct from the other two knockout cell types. To increase statistical power for identifying non-canonical autophagy-dependent differences in gene expression, and control for differences between knockout cells (which may be due to effects on cell biology unrelated to autophagy or TLR signaling or due off target effects of CRISPR), we focused on genes that showed differences in expression between control cells and combined knockout cell lines.

Based on our hypothesis that non-canonical autophagy contributes to limiting GC B cell response, we first asked whether disruption of autophagy led to increase in expression of genes in response to TLR stimulation. However, comparison of control (parental cells) and *ATG5, ATG7* and *ITGAV*-KO cells did not reveal large differences in transcriptional responses to CpG treatment. Instead many of the TLR-responsive genes from the control cells showed increased expression in unstimulated knockout cells (**Fig 7A,B**). This is shown for *ATG5* knockout cells in figure **7A** and we observed similar pattern in all three knockout cells **(Fig 7B**). Many of these genes continued to exhibit upregulation upon stimulation in the knockout cells, but this induction was generally not more pronounced than in the parental cells. Genes that exhibited this pattern included genes related to B cell activation (*AICDA*), TLR signaling (*NFKB1, RELA)*, putative autophagy-regulated genes from the primary B cell analysis (*BCL21L1, TNF, and MYBBP1A*), as well as other genes that have not been well characterized in B cells (such *as SIRPA* and *RASGRP1*), (**Fig 7B and sup Fig 3**). Thus, while non-canonical autophagy is involved in limiting expression of TLR-responsive genes, a major role for non-canonical autophagy may be in regulating basal state of such genes. We therefore focused majority of our analysis on unstimulated conditions, again focusing on changes common to all three knockouts.

**Fig 7:**
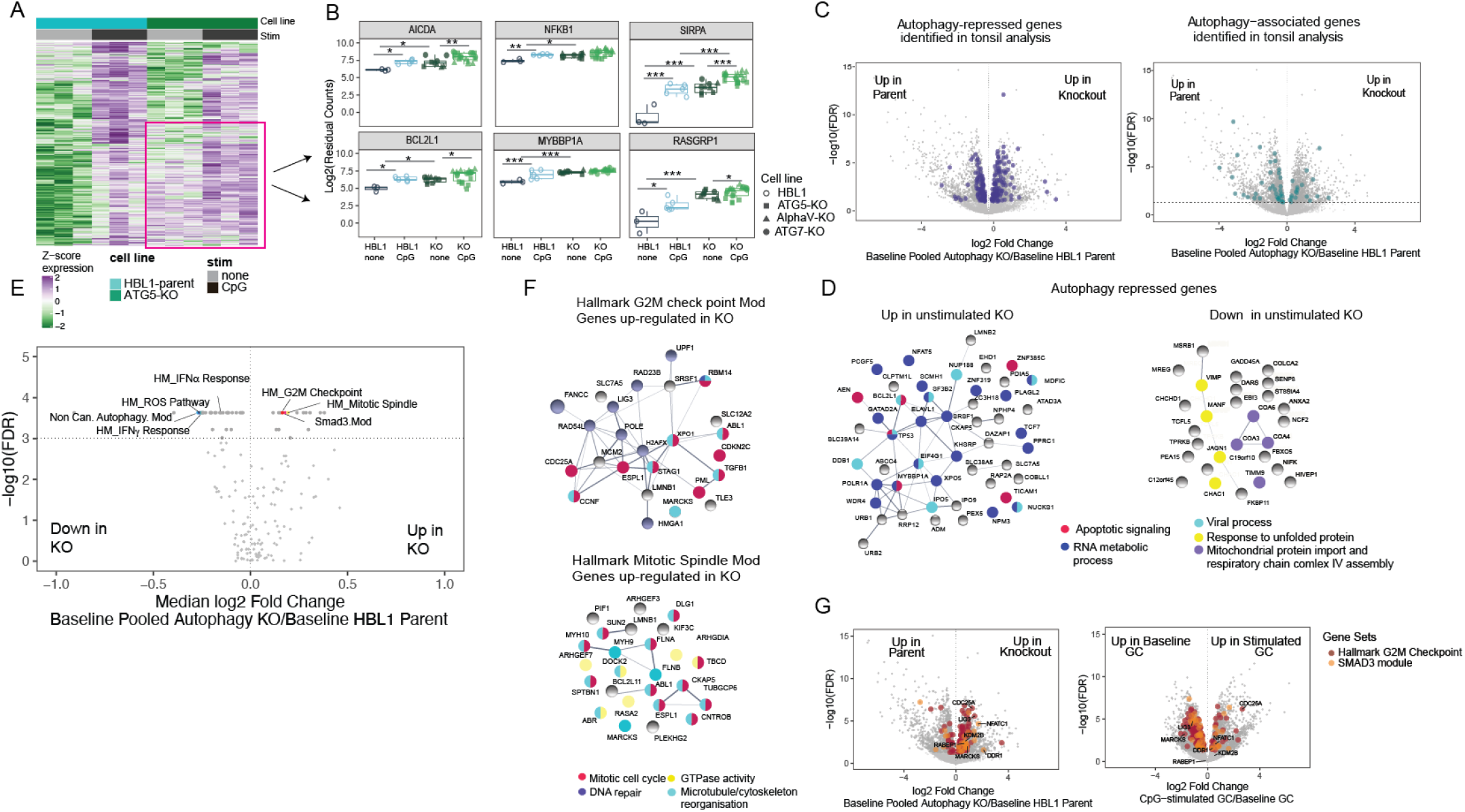
Non-canonical autophagy limits TLR responses through cell cycle related genes: HBL1 parental cell line and cell lines with *ATG5*-CRISPR, *ITGA*V-CRISPR or *ATG7-*CRISPR were stimulated with either CpG 2395 or 2006 for 3h or left unstimulated and subjected to RNA-seq. Data are all from 3 technical replicates for each condition: unstimulated or stimulated for each cell line, used for RNA-sequencing experiment. Stimulation data includes samples stimulated with CpG 2395 and CpG 2006. **(A)** Heat map of genes induced upon stimulation in the parental cells and /or the ATG5 knockout cells. Genes were hierarchically clustered using Euclidean distance & complete linkage; colors represent scaled gene expression values. Data generated from three replicates of unstimulated or stimulated parental cells or knockout cells. **(B)** Individual gene plots showing batch corrected and normalized log2 counts for gene expression. Each dot represents data from a single sample from each cell line and data from n=3 experiments are plotted *p*-values are based on multiple t tests corrected for using Benjamini and Hochberg’s methods and presented as * (*p*<0.05) or ** (*p*<0.01) ***(*p*<0.001). **(C)** Volcano plots showing different expressed genes between combined unstimulated parental cells and combined knockout cells (pooled knockout. Highlighted are genes from the autophagy associated or autophagy regulated sets from the tonsil analysis that also showed significant changes in the combined unstimulated knockout cells. **(D)** Genes significantly up-regulated or down-regulated by 1.5 fold in the unstimulated knockout cells from the autophagy regulated set are plotted as a network, and the known function of these genes are highlighted based on the string pathway analysis tool. (**E**) Volcano plot showing changes in gene module expression between combined unstimulated parental lines and combined unstimulated knockout cell lines. Dashed line represents significance threshold FDR < 0.001; modules of interest are highlighted. **(F)** Genes significantly upregulated (≥ 1.5-fold) in the unstimulated knockout cells from G2M checkpoint module and mitotic spindle gene set enrichment analysis modules are plotted as a network, and the known function of these genes are highlighted based on the string pathway analysis tool. **(G)** Volcano plots showing differentially expressed genes between combined unstimulated parental cells and combined knockout cells (pooled knockout) or the genes differentially expressed in GC B cells upon stimulation compared to unstimulated cells. Highlighted are the genes from the G2M checkpoint module and the SMAD3 module. Data are representative of replicates from each cell line or 5 different donor tonsil samples.

We next analyzed expression of the putative ‘*autophagy repressed’* genes identified in primary GC B cells (**Fig 7C**). 49 of these genes were upregulated more than 1.5-fold in *ATG5, ATG7* and *ITGAV*-KO cells compared with parental cells, consistent with their repression by autophagy (**Fig 7D**). Furthermore, many were associated with the same cellular processes, including RNA metabolism and apoptotic signaling. In contrast, 28 of these genes were downregulated 1.5-fold in autophagy knockout cells, but fewer of these genes showed association with cellular process, confirming autophagy limits specific pathways (**Fig 7D**). We also analyzed expression of the potential ‘*autophagy associated’* gene set (**Fig 7C**) and found that 21 genes showed 1.5-fold reduced expression in autophagy-knockouts, whereas 10 showed increased expression, confirming that the 21 genes are the autophagy associated genes.

Although analyses of individual genes supported a role for non-canonical autophagy in regulating gene expression, many of the possible autophagy-regulated genes identified in primary B cells were not strongly expressed in HBL-1 cells. To better identify corresponding patterns of gene regulation between these cell systems and hence processes regulated by autophagy in primary B cells, we analyzed gene modules that were differentially expressed in unstimulated autophagy knockouts and unstimulated parental HBL-1 cells (**SupTable2, 7E**). Several gene modules related to cell proliferation and growth were upregulated in autophagy knockout HBL-1 cells, including the Hallmark G2M checkpoint and mitotic spindle modules, which had both shown basal upregulation in the GC cells and upregulated genes from both modules included genes involved in mitosis and microtubule/cytoskeleton re-organization **(Fig 7F**). SMAD3 gene module was another module upregulated in the knockout cells that included genes involved in proliferation (**Fig 7E and G**). Both SMAD3 and G2M modules were downregulated after stimulation in primary GC B cells, consistent with their downregulation by autophagy (**Fig 7G**). In addition, some gene modules identified as potentially autophagy regulated in primary B cells were upregulated in autophagy knockout cells after TLR stimulation, including Myc target genes module which was upregulated in *ATG7-*KO and *ITGAV*-KO cells and the Unfolded Protein Response module was only upregulated in the *ATG7*-KO cells but not in *ITGAV*-KO cells indicating these genes are regulated by macro-autophagy but not by non-canonical autophagy (**Sup Fig 3**). Together, these data indicate that non-canonical autophagy limits activation of genes involved in GC cell growth, proliferation and apoptosis in B cells.

Several gene modules were significantly downregulated in autophagy-knockout cells **(Sup Table2 and Fig 7E)**. Notably, these included type I and type-II IFN response modules, and significantly downregulated genes within these are consistent with a requirement for non-canonical autophagy in activation of IRF7, as well as antigen presentation (**Sup Fig 3**). Curiously, the non-canonical autophagy and Reactive Oxygen Species [ROS] modules, which are related to the mitochondrial and respiratory burst genes upregulated in primary GC cells were also downregulated in autophagy-knockout HBL1 cells (**Sup Fig 3D**). Hence, these data support a role for non-canonical autophagy in promoting type I IFN signaling, antigen presentation and raise the possibility that there is positive feedback in which the transcription of genes involved in autophagy are themselves activated by this process.

## Discussion

Through biochemical and transcriptional analysis of primary B cell populations, here we have shown that human memory and GC B cells acquire the ability to activate autophagy after stimulation. This is associated with increased expression of genes involved in non-canonical autophagy and mitophagy, and increased activation of integrin αvβ3. Complementary experiments in the genetically tractable B cell line HBL-1 demonstrate that autophagy in human B cells is activated by αv integrin and occurs through a mitophagy-related pathway. Furthermore, we show that this non-canonical autophagy pathway regulates TLR signaling and transcriptional responses, limiting expression of genes related to cell proliferation and apoptosis while promoting expression of genes related to antigen presentation and Interferon signaling. These data therefore provide important insights into the complex role of autophagy proteins in human B cell responses.

Human memory and GC B cells were the main B cell sub-populations to show LC3 lipidation in response to TLR stimulation. These data are in agreement with our studies of mouse B cells, and studies from others, which showed that activated mouse B cell populations including marginal zone (MZ) and GC B cells, preferentially activate LC3 after stimulation (13, 14, 16). We also observed low levels of autophagy induction in transitional B cells. Again, this is consistent with our findings in mouse spleen B cells and suggest that B cells utilize this pathway during development and maturation in the bone marrow, but it is downregulated as they become mature naïve B cells. In mouse B cells, we have shown that αvβ3 promotes LC3 recruitment to TLR-containing endosomes, through a non-canonical autophagy pathway involving ROS (14). Autophagy in human GC B cells also requires Integrin αvβ3 and is associated with increased expression of genes associated with non-canonical autophagy and ROS production, indicating the same non-canonical autophagy mechanism in humans. Furthermore, we identified a key role for components of the mitophagy pathway in LC3 lipidation in human B cells. Genes involved in mitochondrial metabolism were also upregulated in GC B cells and we concluded that this increased expression of non-canonical autophagy and mitochondrial modules in GC cells conferred the capacity to activate the autophagy pathway. Many of these genes were also upregulated in naïve cells after stimulation, although this did not appear to be sufficient to allow activation of non-canonical autophagy in naïve cells in our hands. Mitochondrial pathways have been linked with autophagy in other recent studies. Notably, the switch to non-canonical autophagy in mouse GC B cells is linked with changes in mitochondrial homeostasis and requires the gene WIP-2 (13), and mitochondrial proteins have been shown to be prominent in activated B cells from mice (34, 35). Hence, these data highlight an intimate connection between activation induced autophagy and mitochondrial proteins in B cells, that could be explored further in disease settings.

We have previously shown that non-canonical autophagy regulates B cell responses by promoting endolysosomal trafficking of TLRs, which restricts signaling through NF-κB but promotes activation of IRF7. A prediction from this model is that non-canonical autophagy will reduce expression of majority of TLR-response genes (those activated by NF-κB) but will activate the expression of a new subset of genes. Comparison of the transcriptome of naïve and GC B cells after stimulation follow this prediction, allowing identification of potential autophagy repressed and activated genes, and further supporting our model, many of the autophagy-repressed genes correspond to known NF-κB targets. GC B cells also showed increased expression of reported IRF7 gene signatures compared with naïve cells, in agreement with our model. However, we did not observe strong activation of these genes after stimulation, and the autophagy activated genes did not correspond closely with known IRF7-activated genes. We hypothesize that this reflects cell-specific differences in IRF7 target genes (many of the published gene sets are derived from DCs or other immune cells) and timing of gene expression measurement. However, it is also possible that autophagy promotes activation of additional IRFs or other transcription factors that are responsible for activation of these genes.

Analysis of differentially expressed gene modules revealed cellular processes that were differentially activated in naïve and GC B cells, and comparison with results from autophagy-knockout HBL-1 cells provided supporting evidence for a role of autophagy in regulating several of these. In particular non-canonical autophagy reduced activation of genes involved in microtubule re-organization, cell division, apoptosis and transcriptional regulation, suggesting that autophagy regulates the ability of GC B cells to divide and proliferate following antigenic stimulation. Conversely, non-canonical autophagy promoted expression of genes involved in antigen presentation, cytokine signaling and IFN signaling, consistent with the role of autophagy in activation of IRF transcription factors and type I IFN production (17). Autophagy-knockout HBL-1 cells also provided additional insights into the role of non-canonical autophagy in GC B cell function. First, we identified additional autophagy regulated genes. Notably genes involved in somatic hypermutation such as *AICDA*, and TLR signaling genes were upregulated in knockout cells, in agreement with our findings in B cell-specific αv-knockout mice (16), and we also observed increased expression of other signaling genes including *SIRPA, RASGRP1* and *MYBBP1A* which have not been studied in B cells. Second, it was notable that the major effect of deletion of autophagy components was to increase expression of TLR-responsive genes in unstimulated cells, rather than to increase their activation after treatment with exogenous TLR ligands, and this was accompanied by increased NF-κB activation. We speculate that a major role for αv-induced non-canonical autophagy may be to suppress this basal activation of TLR signaling in B cells that are poised to respond to antigens including self-reactive B cells. HBL1 cells harbor mutations that can cause activation of MyD88 and the BCR and TLR9 are already colocalized in these cells in the absence of exogenous ligands (31). Therefore, this cell line may be particularly sensitive to this mechanism of basal regulation. However, similar low level basal activation of self-reactive B cells has also been reported in culture, probably due to activation by cellular debris containing endogenous TLR ligands (36). Furthermore, we consistently observe basal NF-κB activation and proliferation in B cells from αv and Atg5-knockout mice, both in culture and directly after isolation, supporting the idea that non-canonical autophagy regulates basal TLR activation in primary non-transformed cells (14, 16). Moreover, basal activation of TLR pathway has been reported in immune cells from SLE subjects and another family of integrin has also been implicated in basal suppression of TLR signaling (37, 38). Thus, loss of this basal suppression mechanism could contribute to autoimmunity, by lowering the threshold for self-reactive B cells to escape tolerance or undergo class-switching.

HBL-1 cells are activated B cell like (ABC) diffuse large B cell lymphoma (DLBCL) cell line that arise from post germinal center B cells (39) and retain some properties of these cells. Relevant to our studies they have been shown to undergo self-antigen induced BCR signaling and grow in the presence of cellular debris (40). Despite the caveat of high basal NF-κB activation in these cells, robust induction of NF-κB and lipidation of LC3 upon stimulation, indicated that they were appropriate for our experiments. Furthermore, their growth in cell debris proved to be particularly advantageous for highlighting mechanisms of regulation of self-reactive B cells. A significant overlap in gene expression changes in *ITGAV*-KO cells with *ATG7*-KO cells further established αv as an important component of the non-canonical autophagy pathway. However, we were surprised to find significant differences in the *ATG5* and *ATG7*-KO cells from the gene expression analysis (**Fig 7, Sup Fig 3A**). It remains to be determined whether this is an effect seen only in the cell line or if it is relevant for specific function of ATG5 and ATG7.

Genetic variants in autophagy genes, including *ATG5*, are associated with increased risk of systemic lupus erythematosus (SLE) (8, 41) and disruption of autophagy has been reported in a range of autoimmune diseases (42, 43). However, the complex role of autophagy in B cells and other immune cell types have made it difficult to identify the mechanisms by which autophagy may protect or contribute to disease. Our data highlight the advantages of studying specific autophagy pathways in a single cell type, and across distinct cell populations in understanding this complex problem. In B cell-focused experiments in mice, we have previously shown that non-canonical autophagy plays a critical role in regulating autoreactive responses, and disruption of non-canonical autophagy only in B cells is sufficient to increase generation of autoantibodies and accelerate lupus-like autoimmunity (14, 15). The confirmation that this non-canonical pathway is active in human B cells, and regulates transcriptional responses to stimulation, provides a potential mechanism by which loss of ATG5 function and altered non-canonical autophagy may promote SLE. Although, the full contribution of macro-autophagy and non-canonical autophagy to autoimmunity is likely to be complex, our data specify pathways that can be investigated to unravel such complexities in terms of B cell dysregulation during development of lupus.

## Materials and Methods

### Purification of B lymphocytes from human tissue

Tonsils were obtained from patients undergoing tonsillectomies and spleens were obtained from patients undergoing pancreatectomy in accordance with an IRB approved protocol. Tissues were harvested and processed in RPMI (10% fetal bovine serum, 2mM glutamine, 100U/ml penicillin and 100μg/ml streptomycin, and 50mM 2-b-mercaptoethanol) to generate single cell suspensions which were stored as frozen aliquots. B cells were isolated from single cell suspensions by using negative selection kit (Stem Cell Technologies) prior to flow cytometry and cell sorting.

### Antibodies, reagents, and cell lines

Anti-human antibodies used for flow cytometry include antibodies against: CD19-PEcy7 (HIB19; BD Biosciences), CD24-BV421 (ML5; BioLegend), CD10-APC (HI10; BioLegend), CD27-BV605 (L128; BD Biosciences), CD38-PerCP-Cy5.5 (HIT2, BD Biosciences), IgD-BV510 (IA6-2; BD Biosciences), CD51-PE (NKI-M9; BioLegend), IgG2a, κ Isotype Control-PE (G155-178; BD Biosciences). Type C CPG ODN 2395, Type B CPG ODN 2006 and R848 were from Invivogen. Antibodies anti-NF-κB p65 (D14E12), anti-LSD1 (C69G12), anti-Histone (D1H2) and horseradish peroxidase conjugated anti-rabbit IgG were from Cell Signaling Technology. Anti-LC3B and anti-actin (AC-74) were from Sigma-Aldrich. Anti-p62 antibody was from American Research Products. Anti-IRF7 antibodies (H-246) and (AHP1180) were from Santa Cruz Biotechnology Inc and Bio-Rad Laboratories Inc, respectively. Anti-IgM was from Jackson Immuno Research Laboratories and sCD40 ligand was from Peprotech. The human diffuse large B cell lymphoma (HBL1) cell line was provided by Dr. Richard James (James Lab, Seattle Children’s Research Institute, Seattle, WA).

### Flow cytometry and cell sorting

Frozen aliquots of single cell suspension from tonsils or spleens were harvested in PBS/ 0.5% BSA/ 2 mM EDTA. Single cell suspensions were blocked with Fc Block (Biolegend) and stained with fluorochrome tagged antibodies for surface markers at 4°C for 30 min. Samples were acquired using LSRII flow cytometer (Becton and Dickinson) and analyzed by FlowJo software (Tree Star Inc.). For sorting of tonsil or spleen B cell populations, after B cell enrichment with the negative selection cocktail (Stem Cell Technologies), cells were labeled with anti-CD19-PEcy7, anti-CD24-BV421, anti-CD10-APC, anti-CD27-BV605, anti-CD38-PerCPcy5.5, anti-IgD-BV510 antibodies then sorted at 4°C with FACS Aria (BD Bioscience). Post-sort purity was checked, and cells were rested for 1 hr at 37°C with 5% CO_2_ in RPMI (10% fetal bovine serum, 2mM glutamine, 100U/ml penicillin and 100μg/ml streptomycin, and 50mM 2-b-mercaptoethanol) before being used for stimulation or RNA extraction. For expansion of clones, transfected HBL1 cell mixes were single cell sorted into multiple 96 well plates containing 200μl of complete IMDM, GlutaMAX media supplemented with 10% fetal bovine serum, 50U/ml penicillin and 50μg/ml streptomycin, and 50μM 2-β-mercaptoethanol and cultured at 37°C with 5% CO_2_.

### RNA sequencing

#### Bulk RNA-sequencing

The indicated populations were isolated by flow cytometry, and aliquots were sorted directly into SMARTer v4 lysis reagents (Clontech). Cells were lysed and cDNA was synthesized. After amplification, sequencing libraries were prepared using the Nextera XT DNA Library Preparation Kit (Illumina). Barcoded libraries were pooled and quantified using a Qubit® Fluorometer (Life Technologies). Single-read sequencing of the pooled libraries was carried out on a HiSeq2500 sequencer (Illumina) for 74 cycles, using TruSeq v3, and SBS kits (Illumina). Target read depths were ∼5-10 million raw reads per sample.

#### RNA-sequencing pipeline analysis

Raw RNA-sequencing reads were processed using a local Galaxy server (46, 47). A minimum quality score of 30 was enforced for all reads by trimming bases from 5’ and 3’ ends using the FASTQ Quality Trimmer tool in Galaxy. Trimmed reads were then aligned to the GRCh38 (release 77) reference genome using STAR. Raw gene counts were generated using htseq-count (v.0.5.4p3) (49), based on the union of exons for each gene as described in the Ensembl gene model GTF file for GRCh38. Quality metrics for aligned reads were obtained using the Picard (v.1.56) suite of tools (http://picard.sourceforge.net). To ensure the highest data quality, we sequentially examined distributions of un-related Picard quality control metrics (PF_ALIGNED_BASES and MEDIAN_CV_COVERAGE) and eliminated outlier libraries. Counts were normalized using the trimmed mean of M values (TMM). Genes were included in analyses if they had greater than 1 count per million in at least 10% of libraries. Differential expression of individual genes was determined with limma-voom. Study subject ID was used as a blocking variable when comparing multiple treatments or cell subsets taken from the same individuals. Raw *P* values were corrected for multiple testing. Gene set analyses were performed using mroast; determined PPI interactions using STRING.

### Western Blots

Nuclear and cytoplasmic extracts were prepared by lysing cells in hypotonic nuclear extraction buffer (1M Hepes, pH7.5, 5M NaCl, 0.5M EDTA pH8, 50% glycerol, 10% Igepal, 10% TritonX100) supplemented with protease inhibitor cocktail (Pierce) for 10 min followed by centrifugation at 1500g for 5 min at 4°C to pellet the nuclei and nuclei were resuspended in RIPA buffer. Lysates were centrifuged for 10 min at 4°C at 14000g and supernatant was collected as nuclear fraction. separated by electrophoresis using NuPage-Bis-Tris gels (Invitrogen) and blotted onto PVDF membranes. Non-specific binding was blocked with 5% BSA in TBS-Tween (0.1%) followed by incubation with primary antibodies (1:1000 dilution) overnight at 4°C and secondary antibody horseradish peroxidase conjugated antibodies (1:5000 dilution) for 1 hr at room temperature. Membranes were washed thoroughly with TBS-Tween (0.1%) after antibody incubations and developed using ECL reagents (Millipore). For re-probing blots were stripped for 30 min at 37 °C with Restore PLUS stripping buffer (Thermo Scientific).

### CRISPR targeting

CRISPR crRNAs targeting BNIP3, ATG5, ATG7, and integrin αv were identified using the Broad Institute single guide RNA design tool and synthesized (IDT) containing phosphorothioate linkages and 2’ O methyl modifications. crRNAs and trans-activating crRNA (tracrRNA; IDT) hybrids were mixed with Cas9 nuclease (IDT) at the ratio of 1.2:1 and transfected into cells by Neon electroporation (Thermo Fisher Scientific). Transfected cell mixture was single cell sorted and total genomic DNA was extracted from the single cell clones using *Quick*-DNA Microprep Kit (ZymoResearch). To assess knockout efficiency, guide target genomic regions were amplified using PrimeSTAR GXL DNA Polymerase (Takara Bio) with primers about 250 – 350 bp away from the guide target site. PCR products were purified using GeneJet PCR Purification Kit (Thermo Fisher Scientific) and analyzed by Sanger Sequencing and Inference of CRISPR Edits (ICE; *ice.synthego.com*). Flow cytometry or western blot was used to validate knockdown on a protein level.

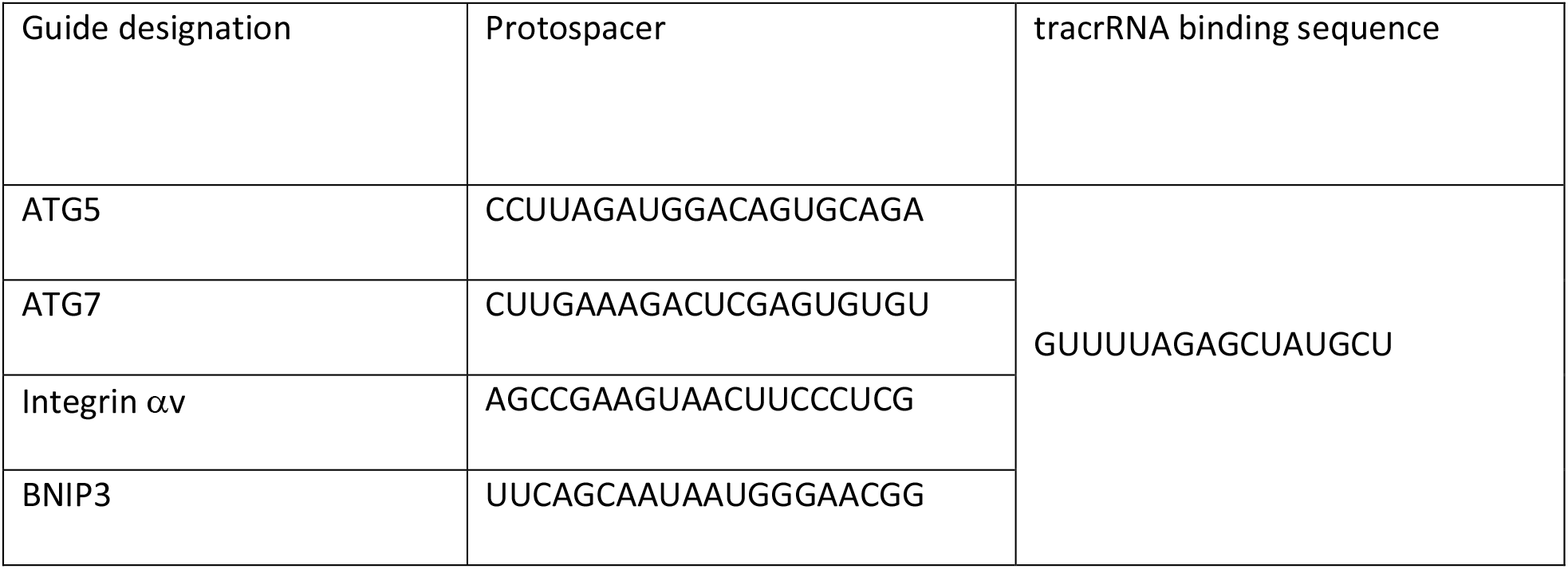

### Super-resolution microscopy

Super-resolution microscopy on tonsil B cell subsets was performed as previously described(14, 44). Briefly, FACs sorted tonsil B cell subsets were seeded on to poly-L-lysine coated chamber slides (Nunc Lab-Tek, Thermo Scientific) stimulated, fixed and stained as previously described (14). Integrin staining was performed using primary αvβ3 (23C6) or LIBS αvβ3 (22703) antibody and anti-rabbit atto-488 (Hypermol, Bielefeld, Germany) or streptavidin 647 as secondary reagents. Samples were imaged at room temperature using a customized Nikon N-STORM with 60X oil immersion and a “Perfect Focus” system. Imaging was performed in an extracellular solution containing reducing and oxygen scavengers, as specified by dSTORM protocols(45). After activation the fluorochromes were converted into a desired density of single molecules per frame and imaged continuously at 10,000–30,000 frames. Localization was reconstructed using Nikon Elements Imaging software.

## Acknowledgements

We thank all members of the Acharya, Lacy-Hulbert and James’s laboratories for assistance and advice on this project. We thank the flow cytometry and genomics core facilities at Benaroya Research Institute for expert assistance and Dr. Simon Goodman at Merk laboratories for integrin antibodies.

## Author contributions

Data analysis and experimental work: MA, SS, VM.

Additional assistance in experimental design, execution and analysis: RGJ, ES and IM (CRISPR/Cas9), JMT, ECG and CVM (STORM imaging), NG (tonsil cell subpopulation analysis) and DS (human spleen samples Conceptualization: MA, and ALH.

Writing: ALH, SS, VM and MA

## Data and materials availability

All data from sequencing will be made publicly available.

