## supplemental materials for "Transcriptomic analysis of pathways associated with αv integrin-related non-canonical autophagy in human B cells"

### This PDF file includes:

Supplementary Text
Figs. S1 to S3
Tables S1 to S2

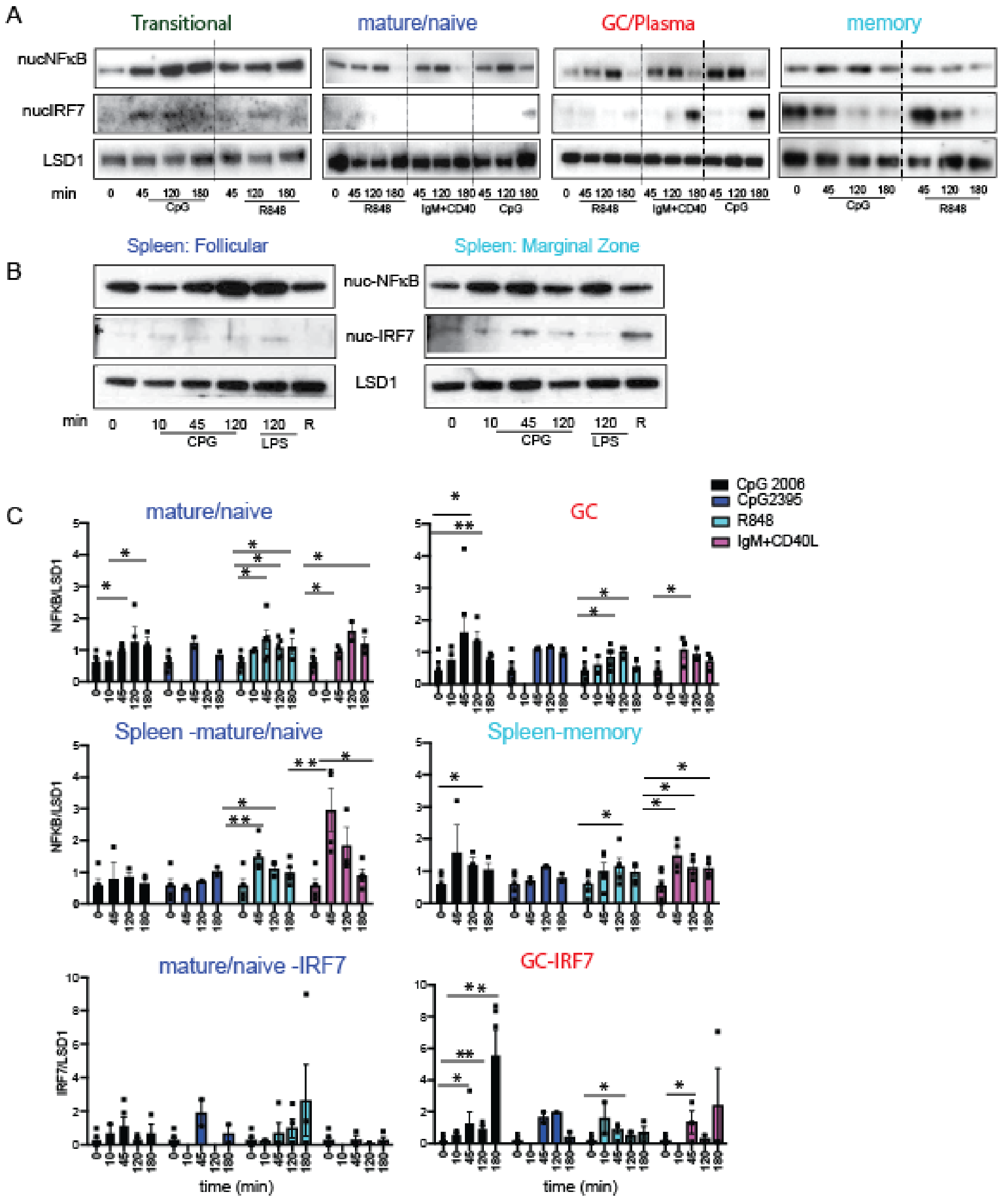


Supplementary Fig 1: Activation induced NFκB/IRF7 signaling in human B cell subsets

**Supplementary Fig 1: Response to stimulation in tonsil B cell subsets: (A)** Western blot analysis for LC3 expression and processing in sorted B cell subsets (200,000) from human tonsils. Sorted B cells were stimulated for the indicated times with CpG DNA or R848 (TLR ligand stimulation or anti-IgM+ CD40L (non-TLR ligand) for indicated time (mins) and nuclear extracts from cell lysates were used to assess NF-kB or IRF7 activation (translocation to the nucleus). LSD1 is shown as nuclear loading control. **(B)** Western blot analysis of NF-kB or IRF7 activation from sorted spleen follicular or marginal zone B cell subsets from nuclear extracts after stimulation for indicated times with CpG DNA or R848 or LPS (10mg/mL) (TLR ligand stimulation). **(C)** Quantification of NF-kB or IRF7 activation from A and B by densitometry. Western blots are representative blots from 8 different donors and quantification is based on data from 3-8 donors.

A

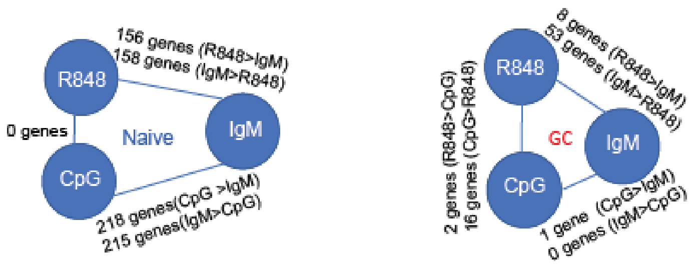


B

Modules changing in similar direction in both subsets upon stimulation

| | |
|---|---|
| HM_ Myc Targets V2<br>HDAC2.MOD | Higher basal expression in GC cells compared to Naive.<br>Significant Up-regualtion in both subsets but, extent of upregulation : Naive cells > GC cells. |
| HM_Myc Target V1<br>IL27RA.MOD<br>HM_Unfolded Protein Resp.Mod<br>TFAM.Mod | Higher Basal expression in GC cells.<br>Significant Up-regualtion in Naive cells and trends towards upregulation in GC cells, extent of upregulation: Naive cells >GC cells |
| Go Endosomes.Mod | Lower basal expression GC cells.<br>Significant up-regulation upon stimulation in both subsets. |
| CD19.Mod<br>IRF8. Mod<br>HM_ Interferon-a Resp.Mod<br>SMAD3<br>NCOA3 | Lower basal expression in GC compared to Naive.Significant down-regulation in both subsets but extent of downregulation: Naive cells > GC cells. |
| Stat1. Mod<br>Stat2.Mod<br>HM_ Interferon-γ Resp.Mod<br>IRF7. Mod | Lower basal expression in GC cells.<br>Significant down-regulation in Naive cells and only a trend towards down-regulation in GC cells. |

Modules changes only in the GC cells and not in Naive cells

| | |
|---|---|
| E2F target.Mod<br>Hallmark G2M checkpoint | Higher basal expression in GC cells.<br>Significant down-regulation in GC cells and only a trend towards down-regualtion in Naive cells |
| Stat3.Mod<br>Stat6.Mod<br>SP1.Mod | Trend towards up-regulation in GC cells but trend towards down-regulation in Naive cells (Change upon stimulation not significant in both subsets with any stimulation). |

Supplementary Fig 2 : Differentially expressed genes and modules between stimulations in naive and GC B cells.

**Supp. Fig 2: Differentially expressed genes in Naïve and GC B cell subsets: (A)** Schematic of differentially expressed (DE) genes from pairwise comparisons between each of the three stimulation CpG, R848 and anti-IgM+CD40L in naïve and GC B cells. Numbers of significantly DE genes for each comparison are shown. **(B)** Table for module changes in GC and naïve cells based on Roast analysis. Modules showing significant changes in Naïve cells with at least two different stimulation condition and showing a similar trend in the GC cells are represented

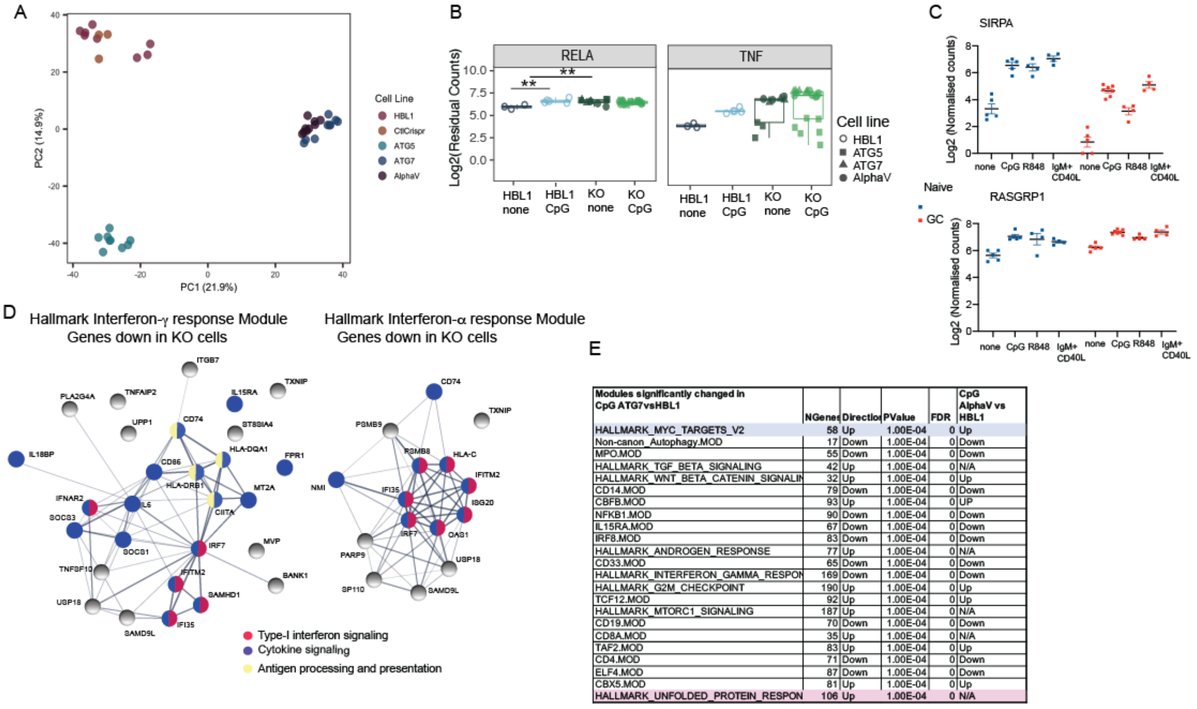


| Modules significantly changed in CpG ATG7vsHBL1 | NGenes | Directio | PValue | FDR | CpG AlphaV vs HBL1 |
|---|---|---|---|---|---|
| HALLMARK_MYC_TARGETS_V2 | 58 | Up | 1.00E-04 | 0 | Up |
| Non-canon_Autophagy.MOD | 17 | Down | 1.00E-04 | 0 | Down |
| MPO.MOD | 55 | Down | 1.00E-04 | 0 | Down |
| HALLMARK_TGF_BETA_SIGNALING | 42 | Up | 1.00E-04 | 0 | N/A |
| HALLMARK_WNT_BETA_CATENIN_SIGNALIN | 32 | Up | 1.00E-04 | 0 | Up |
| CD14.MOD | 79 | Down | 1.00E-04 | 0 | Down |
| CBFB.MOD | 93 | Up | 1.00E-04 | 0 | UP |
| NFKB1.MOD | 90 | Down | 1.00E-04 | 0 | Down |
| IL15RA.MOD | 67 | Down | 1.00E-04 | 0 | Down |
| IRF8.MOD | 83 | Down | 1.00E-04 | 0 | Down |
| HALLMARK_ANDROGEN_RESPONSE | 77 | Up | 1.00E-04 | 0 | N/A |
| CD33.MOD | 65 | Down | 1.00E-04 | 0 | Down |
| HALLMARK_INTERFERON_GAMMA_RESPON | 169 | Down | 1.00E-04 | 0 | Down |
| HALLMARK_G2M_CHECKPOINT | 190 | Up | 1.00E-04 | 0 | Up |
| TCF12.MOD | 92 | Up | 1.00E-04 | 0 | Up |
| HALLMARK_MTORC1_SIGNALING | 187 | Up | 1.00E-04 | 0 | N/A |
| CD19.MOD | 70 | Down | 1.00E-04 | 0 | Down |
| CD8A.MOD | 35 | Up | 1.00E-04 | 0 | N/A |
| TAF2.MOD | 83 | Up | 1.00E-04 | 0 | Up |
| CD4.MOD | 71 | Down | 1.00E-04 | 0 | Down |
| ELF4.MOD | 87 | Down | 1.00E-04 | 0 | Down |
| CBX5.MOD | 81 | Up | 1.00E-04 | 0 | Up |
| HALLMARK_UNFOLDED_PROTEIN_RESPON | 106 | Up | 1.00E-04 | 0 | N/A |

Supplementary Fig 3: Genes and pathways regulated by non-canonical autophagy

**Supplementary Fig 3: Genes regulated by non-canonical autophagy in human B cells**: **(A)** PCA plot for indicated cell populations. Each dot represents a technical replicate for each cell line. Data represent cells with both stimulated and unstimulated condition. Parental cells with guides but without Cas9 are shown as CRISPR controls. **(B)** Single gene plots, showing batch corrected and normalized log2 counts for gene expression. Each dot represents data from a single sample from each cell line and data from n=3 experiments are plotted. **(C)** Graph shows normalized log2 counts for gene expression of the genes *SIRPA* and *RASGRP1* in sorted B cell subsets with or without stimulation. Each dot represent data from a single donor among the 5 different donors. **(D)** Analysis of network of genes from the mentioned modules that show significant down-regulation in pooled knockout cells, using string pathway analysis tool, the known function of these genes are highlighted based on the string pathway analysis tool. **(E)** Table showing overall module changes in unstimulated parental cells and either *ATG7*-KO cells or *ITGAV*-KO cells using Roast analysis. Highlighted are the two modules that were identified in tonsil GC B cell analysis in **Fig 6A.**

**CpG**

| | | NGenes | Direction | PValue | FDR |
|---|---|---|---|---|---|
| GC | CD19.MOD | 77 | Down | 1.00E-04 | 0.00078629 |
| | HALLMARK_G2M_CHECKPOINT | 192 | Down | 1.00E-04 | 0.00078629 |
| | IRF8.MOD | 88 | Down | 1.00E-04 | 0.00078629 |
| | NCOA3.MOD | 75 | Down | 1.00E-04 | 0.00078629 |
| | SMAD3.MOD | 92 | Down | 1.00E-04 | 0.00078629 |
| | HALLMARK_INTERFERON_ALPHA_RESPONSE | 93 | Down | 1.00E-04 | 0.00078629 |
| | EOMES.MOD | 75 | Down | 1.00E-04 | 0.00078629 |
| | HALLMARK_E2F_TARGETS | 196 | Down | 5.98E-13 | 1.03E-10 |
| | FOS.MOD | 68 | Down | 3.58E-05 | 0.00088584 |
| | HDAC2.MOD | 89 | Up | 1.00E-04 | 0.00078629 |
| | HALLMARK_MYC_TARGETS_V2 | 58 | Up | 1.00E-04 | 0.00078629 |
| | GO_endosome.MOD | 58 | Up | 1.00E-04 | 0.00078629 |
| Naïve | CD19.MOD | 77 | Down | 1.00E-04 | 0.00021096 |
| | IRF8.MOD | 88 | Down | 1.00E-04 | 0.00021096 |
| | HALLMARK_INTERFERON_ALPHA_RESPONSE | 93 | Down | 1.00E-04 | 0.00021096 |
| | NCOA3.MOD | 75 | Down | 1.00E-04 | 0.00021096 |
| | STAT2.MOD | 89 | Down | 1.00E-04 | 0.00021096 |
| | STAT1.MOD | 92 | Down | 1.00E-04 | 0.00021096 |
| | AFF1.MOD | 85 | Down | 1.00E-04 | 0.00021096 |
| | IRF7.MOD | 90 | Down | 1.00E-04 | 0.00021096 |
| | SMAD3.MOD | 92 | Down | 1.00E-04 | 0.00021096 |
| | HALLMARK_INTERFERON_GAMMA_RESPONSE | 189 | Down | 1.00E-04 | 0.00021096 |
| | NFATC3.MOD | 84 | Down | 1.00E-04 | 0.00021096 |
| | FOS.MOD | 68 | Down | 1.00E-04 | 0.00021096 |
| | PHF1.MOD | 76 | Down | 1.00E-04 | 0.00021096 |
| | SP1.MOD | 97 | Down | 1.00E-04 | 0.00021096 |
| | IKZF1.MOD | 83 | Down | 1.00E-04 | 0.00021096 |
| | FLI1.MOD | 97 | Down | 1.00E-04 | 0.00021096 |
| | TIGIT.MOD | 81 | Down | 1.00E-04 | 0.00021096 |
| | TOX.MOD | 71 | Down | 1.00E-04 | 0.00021096 |
| | RUNX3.MOD | 74 | Down | 1.00E-04 | 0.00021096 |
| | STAT4.MOD | 73 | Down | 1.00E-04 | 0.00021096 |
| | HALLMARK_HYPOXIA | 167 | Down | 1.00E-04 | 0.00021096 |
| | Autophagy_SAB.MOD | 77 | Down | 1.00E-04 | 0.00021096 |
| | HALLMARK_KRAS_SIGNALING_UP | 144 | Down | 1.00E-04 | 0.00021096 |
| | HALLMARK_P53_PATHWAY | 182 | Down | 1.00E-04 | 0.00021096 |
| | TCF7.MOD | 66 | Down | 1.00E-04 | 0.00021096 |
| | HALLMARK_HEME_METABOLISM | 171 | Down | 1.00E-04 | 0.00021096 |
| | CHD3.MOD | 69 | Down | 1.00E-04 | 0.00021096 |
| | CD5.MOD | 48 | Down | 1.00E-04 | 0.00021096 |
| | BAZ1A.MOD | 87 | Down | 0.00019998 | 0.00054057 |
| | Kegg_lysosome.MOD | 112 | Down | 0.00019998 | 0.00054057 |
| | MPO.MOD | 59 | Down | 0.00019998 | 0.00054057 |
| | GZMM.MOD | 77 | Down | 0.00019998 | 0.00054057 |
| | CD2.MOD | 57 | Down | 0.00019998 | 0.00054057 |
| | HALLMARK_KRAS_SIGNALING_DN | 110 | Down | 0.00019998 | 0.00054057 |
| | KLRD1.MOD | 61 | Down | 0.00019998 | 0.00054057 |
| | HALLMARK_COMPLEMENT | 160 | Down | 0.00029997 | 0.00081596 |
| | HALLMARK_BILE_ACID_METABOLISM | 92 | Down | 0.00029997 | 0.00081596 |
| | CD28.MOD | 61 | Down | 0.00029997 | 0.00081596 |
| | HALLMARK_MYC_TARGETS_V2 | 58 | Up | 1.00E-04 | 0.00021096 |
| | HDAC2.MOD | 89 | Up | 1.00E-04 | 0.00021096 |
| | IL27RA.MOD | 93 | Up | 1.00E-04 | 0.00021096 |
| | HALLMARK_MYC_TARGETS_V1 | 197 | Up | 1.00E-04 | 0.00021096 |
| | TAF9.MOD | 95 | Up | 1.00E-04 | 0.00021096 |
| | TFAM.MOD | 70 | Up | 1.00E-04 | 0.00021096 |
| | PATZ1.MOD | 85 | Up | 1.00E-04 | 0.00021096 |
| | MTA1.MOD | 92 | Up | 1.00E-04 | 0.00021096 |
| | HALLMARK_UNFOLDED_PROTEIN_RESPONSE | 106 | Up | 1.00E-04 | 0.00021096 |
| | CBX5.MOD | 82 | Up | 1.00E-04 | 0.00021096 |
| | GO_endosome.MOD | 58 | Up | 1.00E-04 | 0.00021096 |
| | YY1.MOD | 86 | Up | 1.00E-04 | 0.00021096 |
| | HALLMARK_REACTIVE_OXIGEN_SPECIES_PATHWAY | 45 | Up | 0.00029997 | 0.00081596 |
| | Modified_biocarta_NFKB.MOD | 27 | Up | 0.00029997 | 0.00081596 |

**IgM+CD40**

| | | NGenes | Direction | PValue | FDR |
|---|---|---|---|---|---|
| GC | HALLMARK_E2F_TARGETS | 196 | Down | 9.53E-16 | 1.65E-13 |
| | CD19.MOD | 77 | Down | 8.64E-11 | 7.48E-09 |
| | HALLMARK_G2M_CHECKPOINT | 192 | Down | 1.75E-08 | 1.01E-06 |
| | HALLMARK_MYC_TARGETS_V2 | 58 | Up | 9.18E-07 | 3.18E-05 |
| Naïve | CD19.MOD | 77 | Down | 1.00E-04 | 0.00050877 |
| | NCOA3.MOD | 75 | Down | 1.00E-04 | 0.00050877 |
| | IRF8.MOD | 88 | Down | 1.00E-04 | 0.00050877 |
| | STAT2.MOD | 89 | Down | 1.00E-04 | 0.00050877 |
| | HALLMARK_INTERFERON_GAMMA_RESPONSE | 189 | Down | 1.00E-04 | 0.00050877 |
| | IKZF1.MOD | 83 | Down | 1.00E-04 | 0.00050877 |
| | AFF1.MOD | 85 | Down | 5.10E-08 | 8.62E-07 |
| | STAT1.MOD | 92 | Down | 1.37E-07 | 1.94E-06 |
| | HALLMARK_INTERFERON_ALPHA_RESPONSE | 93 | Down | 2.06E-07 | 2.68E-06 |
| | IRF7.MOD | 90 | Down | 5.01E-06 | 5.64E-05 |
| | CCNT2.MOD | 85 | Down | 8.78E-05 | 0.00074202 |
| | FLI1.MOD | 97 | Up | 1.00E-04 | 0.00050877 |
| | HALLMARK_MYC_TARGETS_V2 | 58 | Up | 1.00E-04 | 0.00050877 |
| | IL27RA.MOD | 93 | Up | 1.00E-04 | 0.00050877 |
| | HALLMARK_MYC_TARGETS_V1 | 197 | Up | 1.00E-04 | 0.00050877 |
| | TAF9.MOD | 95 | Up | 1.00E-04 | 0.00050877 |
| | TFAM.MOD | 70 | Up | 1.00E-04 | 0.00050877 |
| | HALLMARK_REACTIVE_OXIGEN_SPECIES_PATHWAY | 45 | Up | 1.00E-04 | 0.00050877 |
| | PATZ1.MOD | 85 | Up | 1.00E-04 | 0.00050877 |
| | MTA1.MOD | 92 | Up | 1.00E-04 | 0.00050877 |
| | HALLMARK_UNFOLDED_PROTEIN_RESPONSE | 106 | Up | 1.00E-04 | 0.00050877 |
| | YY1.MOD | 86 | Up | 1.00E-04 | 0.00050877 |
| | HALLMARK_OXIDATIVE_PHOSPHORYLATION | 195 | Up | 8.34E-05 | 0.00074202 |
| | HALLMARK_MTORC1_SIGNALING | 194 | Up | 0.00010127 | 0.00081496 |

**R848**

| | | NGenes | Direction | PValue | FDR |
|---|---|---|---|---|---|
| GC | HALLMARK_E2F_TARGETS | 196 | Down | 1.01E-16 | 1.75E-14 |
| | HALLMARK_G2M_CHECKPOINT | 192 | Down | 2.57E-09 | 1.65E-07 |
| | CD19.MOD | 77 | Down | 2.87E-09 | 1.65E-07 |
| Naïve | CD19.MOD | 77 | Down | 1.00E-04 | 0.00066532 |
| | IRF8.MOD | 88 | Down | 1.00E-04 | 0.00066532 |
| | SMAD3.MOD | 92 | Down | 1.00E-04 | 0.00066532 |
| | HALLMARK_INTERFERON_GAMMA_RESPONSE | 189 | Down | 1.00E-04 | 0.00066532 |
| | TOX.MOD | 71 | Down | 1.00E-04 | 0.00066532 |
| | TCF7.MOD | 66 | Down | 1.00E-04 | 0.00066532 |
| | NKG7.MOD | 57 | Down | 1.00E-04 | 0.00066532 |
| | NCOA3.MOD | 75 | Down | 1.69E-09 | 4.07E-08 |
| | HALLMARK_INTERFERON_ALPHA_RESPONSE | 93 | Down | 7.57E-07 | 1.28E-05 |
| | STAT2.MOD | 89 | Down | 4.55E-06 | 6.99E-05 |
| | STAT1.MOD | 92 | Down | 7.45E-06 | 0.00010486 |
| | FOS.MOD | 68 | Down | 1.21E-05 | 0.00015774 |
| | AFF1.MOD | 85 | Down | 1.45E-05 | 0.00017462 |
| | HDAC2.MOD | 89 | Up | 1.00E-04 | 0.00066532 |
| | HALLMARK_MYC_TARGETS_V1 | 197 | Up | 1.00E-04 | 0.00066532 |
| | TFAM.MOD | 70 | Up | 1.00E-04 | 0.00066532 |
| | HALLMARK_REACTIVE_OXIGEN_SPECIES_PATHWAY | 45 | Up | 1.00E-04 | 0.00066532 |
| | HALLMARK_UNFOLDED_PROTEIN_RESPONSE | 106 | Up | 1.00E-04 | 0.00066532 |
| | HDAC2.MOD | 89 | Up | 9.79E-14 | 5.52E-12 |
| | IL27RA.MOD | 93 | Up | 5.24E-12 | 1.77E-10 |
| | HALLMARK_MYC_TARGETS_V2 | 58 | Up | 7.89E-12 | 2.22E-10 |
| | TAF9.MOD | 95 | Up | 2.77E-08 | 5.85E-07 |
| | TFAM.MOD | 70 | Up | 2.83E-07 | 5.31E-06 |
| | CBX3.MOD | 87 | Up | 1.74E-05 | 0.00019605 |

**Supplementary Table1:** Table for module changes identified using Roast and Camera gene set testing analysis, in sorted GC and Naïve B cells, with or without stimulation (3h) with either CpG or R848 or anti-IgM +CD40L and used for RNA-sequencing**.** Data from stimulation with either CpG 2395 or CpG 2006 were combined and represented as CpG.

| | NGenes | PropDown | PropUp | Direction | PValue | FDR | PValue.Mixe | FDR.Mixed | Stim in Naive | Stim in GC | GC v Naive |
|---|---|---|---|---|---|---|---|---|---|---|---|
| HALLMARK_G2M_CHECKPOINT | 190 | 0.17894737 | 0.48421053 | Up | 1.00E-04 | 0.0002621 | 1.00E-04 | 1.00E-04 | #N/A | Down | Up |
| FOXJ2.MOD | 81 | 0.16049383 | 0.48148148 | Up | 1.00E-04 | 0.0002621 | 1.00E-04 | 1.00E-04 | #N/A | #N/A | #N/A |
| MLL.MOD | 85 | 0.07058824 | 0.47058824 | Up | 1.00E-04 | 0.0002621 | 1.00E-04 | 1.00E-04 | #N/A | #N/A | #N/A |
| HALLMARK_WNT_BETA_CATENIN_SIGNALING | 32 | 0.0625 | 0.46875 | Up | 1.00E-04 | 0.0002621 | 1.00E-04 | 1.00E-04 | #N/A | #N/A | #N/A |
| SMAD3.MOD | 90 | 0.15555556 | 0.44444444 | Up | 1.00E-04 | 0.0002621 | 1.00E-04 | 1.00E-04 | Down | Down | Down |
| RBBP7.MOD | 91 | 0.15384615 | 0.43956044 | Up | 1.00E-04 | 0.0002621 | 1.00E-04 | 1.00E-04 | #N/A | #N/A | Down |
| HALLMARK_MITOTIC_SPINDLE | 188 | 0.14893617 | 0.43617021 | Up | 1.00E-04 | 0.0002621 | 1.00E-04 | 1.00E-04 | #N/A | #N/A | Up |
| TCF12.MOD | 92 | 0.14130435 | 0.42391304 | Up | 1.00E-04 | 0.0002621 | 1.00E-04 | 1.00E-04 | #N/A | #N/A | #N/A |
| RREB1.MOD | 71 | 0.08450704 | 0.4084507 | Up | 1.00E-04 | 0.0002621 | 1.00E-04 | 1.00E-04 | #N/A | #N/A | #N/A |
| TAF2.MOD | 83 | 0.10843374 | 0.39759036 | Up | 1.00E-04 | 0.0002621 | 1.00E-04 | 1.00E-04 | #N/A | #N/A | #N/A |
| TIGIT.MOD | 69 | 0.17391304 | 0.39130435 | Up | 1.00E-04 | 0.0002621 | 1.00E-04 | 1.00E-04 | Down | #N/A | #N/A |
| CBX5.MOD | 81 | 0.13580247 | 0.35802469 | Up | 1.00E-04 | 0.0002621 | 1.00E-04 | 1.00E-04 | Up | #N/A | Down |
| CD28.MOD | 46 | 0.06521739 | 0.2826087 | Up | 1.00E-04 | 0.0002621 | 0.00159984 | 0.00159984 | Down | #N/A | #N/A |
| MTA1.MOD | 92 | 0.15217391 | 0.38043478 | Up | 0.00019998 | 0.00058971 | 1.00E-04 | 1.00E-04 | Up | #N/A | #N/A |
| FUBP1.MOD | 83 | 0.14457831 | 0.32530121 | Up | 0.00019998 | 0.00058971 | 1.00E-04 | 1.00E-04 | #N/A | #N/A | #N/A |
| CD2.MOD | 46 | 0.17391304 | 0.32608696 | Up | 0.00029997 | 0.00090095 | 1.00E-04 | 1.00E-04 | Down | #N/A | #N/A |
| Non-canon_Autophagy.MOD | 17 | 0.58823529 | 0.05882353 | Down | 1.00E-04 | 0.0002621 | 1.00E-04 | 1.00E-04 | #N/A | #N/A | Up |
| MPO.MOD | 55 | 0.58181818 | 0.10909091 | Down | 1.00E-04 | 0.0002621 | 1.00E-04 | 1.00E-04 | Down | #N/A | #N/A |
| HALLMARK_REACTIVE_OXIGEN_SPECIES_PATHWAY | 44 | 0.54545455 | 0.13636364 | Down | 1.00E-04 | 0.0002621 | 1.00E-04 | 1.00E-04 | Up | #N/A | #N/A |
| CBX3.MOD | 86 | 0.51162791 | 0.1627907 | Down | 1.00E-04 | 0.0002621 | 1.00E-04 | 1.00E-04 | #N/A | #N/A | Up |
| CD14.MOD | 79 | 0.50632911 | 0.17721519 | Down | 1.00E-04 | 0.0002621 | 1.00E-04 | 1.00E-04 | #N/A | #N/A | #N/A |
| HALLMARK_DNA_REPAIR | 133 | 0.47368421 | 0.18796993 | Down | 1.00E-04 | 0.0002621 | 1.00E-04 | 1.00E-04 | #N/A | #N/A | Up |
| SMARCD2.MOD | 87 | 0.47126437 | 0.18390805 | Down | 1.00E-04 | 0.0002621 | 1.00E-04 | 1.00E-04 | #N/A | #N/A | #N/A |
| NFKB1.MOD | 90 | 0.46666667 | 0.16666667 | Down | 1.00E-04 | 0.0002621 | 1.00E-04 | 1.00E-04 | #N/A | #N/A | Up |
| HALLMARK_HYPOXIA | 146 | 0.4520548 | 0.15753425 | Down | 1.00E-04 | 0.0002621 | 1.00E-04 | 1.00E-04 | Down | #N/A | #N/A |
| ZBTB7B.MOD | 84 | 0.44047619 | 0.20238095 | Down | 1.00E-04 | 0.0002621 | 1.00E-04 | 1.00E-04 | #N/A | #N/A | Up |
| LC3_TCGA_POS_COR_0.4.MOD | 82 | 0.42682927 | 0.08536585 | Down | 1.00E-04 | 0.0002621 | 1.00E-04 | 1.00E-04 | #N/A | #N/A | #N/A |
| HALLMARK_INTERFERON_GAMMA_RESPONSE | 169 | 0.4260355 | 0.13609468 | Down | 1.00E-04 | 0.0002621 | 1.00E-04 | 1.00E-04 | Down | #N/A | Down |
| IL15RA.MOD | 67 | 0.40298508 | 0.14925373 | Down | 1.00E-04 | 0.0002621 | 1.00E-04 | 1.00E-04 | #N/A | #N/A | #N/A |
| HALLMARK_ALLOGRAFT_REJECTION | 139 | 0.38848921 | 0.16546763 | Down | 1.00E-04 | 0.0002621 | 1.00E-04 | 1.00E-04 | #N/A | #N/A | #N/A |
| HALLMARK_IL6_JAK_STAT3_SIGNALING | 58 | 0.37931035 | 0.24137931 | Down | 1.00E-04 | 0.0002621 | 1.00E-04 | 1.00E-04 | #N/A | #N/A | Down |
| HALLMARK_GLYCOLYSIS | 152 | 0.375 | 0.23026316 | Down | 1.00E-04 | 0.0002621 | 1.00E-04 | 1.00E-04 | #N/A | #N/A | Up |
| CD4.MOD | 71 | 0.32394366 | 0.14084507 | Down | 1.00E-04 | 0.0002621 | 1.00E-04 | 1.00E-04 | #N/A | #N/A | Up |
| CD33.MOD | 65 | 0.32307692 | 0.10769231 | Down | 1.00E-04 | 0.0002621 | 1.00E-04 | 1.00E-04 | #N/A | #N/A | #N/A |
| HALLMARK_XENOBIOTIC_METABOLISM | 131 | 0.26717557 | 0.18320611 | Down | 1.00E-04 | 0.0002621 | 1.00E-04 | 1.00E-04 | #N/A | #N/A | Up |
| HALLMARK_OXIDATIVE_PHOSPHORYLATION | 194 | 0.58247423 | 0.13402062 | Down | 0.00019998 | 0.00058971 | 1.00E-04 | 1.00E-04 | #N/A | #N/A | Up |
| TAF9.MOD | 95 | 0.4631579 | 0.15789474 | Down | 0.00019998 | 0.00058971 | 1.00E-04 | 1.00E-04 | Up | #N/A | Up |
| KLF1.MOD | 49 | 0.40816327 | 0.24489796 | Down | 0.00019998 | 0.00058971 | 1.00E-04 | 1.00E-04 | #N/A | #N/A | #N/A |
| HALLMARK_FATTY_ACID_METABOLISM | 124 | 0.38709677 | 0.15322581 | Down | 0.00019998 | 0.00058971 | 1.00E-04 | 1.00E-04 | #N/A | #N/A | Up |
| HALLMARK_ADIPOGENESIS | 158 | 0.37341772 | 0.26582279 | Down | 0.00019998 | 0.00058971 | 1.00E-04 | 1.00E-04 | #N/A | #N/A | Up |
| HALLMARK_INFLAMMATORY_RESPONSE | 116 | 0.37068966 | 0.18103448 | Down | 0.00019998 | 0.00058971 | 1.00E-04 | 1.00E-04 | #N/A | #N/A | #N/A |
| HALLMARK_INTERFERON_ALPHA_RESPONSE | 86 | 0.36046512 | 0.12790698 | Down | 0.00019998 | 0.00058971 | 1.00E-04 | 1.00E-04 | Down | Down | Down |
| Kegg_lysosome.MOD | 103 | 0.3592233 | 0.16504854 | Down | 0.00019998 | 0.00058971 | 1.00E-04 | 1.00E-04 | Down | #N/A | Down |
| Autophagy_SAB.MOD | 75 | 0.32 | 0.24 | Down | 0.00019998 | 0.00058971 | 1.00E-04 | 1.00E-04 | Down | #N/A | #N/A |
| HALLMARK_COMPLEMENT | 133 | 0.36090226 | 0.23308271 | Down | 0.00029997 | 0.00090095 | 1.00E-04 | 1.00E-04 | Down | #N/A | #N/A |
| CD19.MOD | 70 | 0.35714286 | 0.17142857 | Down | 0.00029997 | 0.00090095 | 1.00E-04 | 1.00E-04 | Down | Down | Down |
| HALLMARK_COAGULATION | 59 | 0.3559322 | 0.11864407 | Down | 0.00029997 | 0.00090095 | 1.00E-04 | 1.00E-04 | #N/A | #N/A | #N/A |

**Supplementary Table2:** Table showing overall module changes in unstimulated parental cells and the pooled knockout cells, using Roast analysis. Changes in the gene sets related to GC and Naïve cells are also indicated.
